# A conserved BAH module within mammalian BAHD1 connects H3K27me3 to Polycomb gene silencing

**DOI:** 10.1101/2021.03.11.435004

**Authors:** Huitao Fan, Yiran Guo, Yi-Hsuan Tsai, Aaron J. Storey, Arum Kim, Weida Gong, Ricky D. Edmondson, Samuel G. Mackintosh, Haitao Li, Stephanie D. Byrum, Alan J. Tackett, Ling Cai, Gang Greg Wang

**Affiliations:** Lineberger Comprehensive Cancer Center, University of North Carolina at Chapel Hill School of Medicine, Chapel Hill, NC 27599, USA; Department of Biochemistry and Biophysics, University of North Carolina at Chapel Hill School of Medicine, Chapel Hill, NC 27599, USA; Curriculum in Genetics and Molecular Biology, University of North Carolina at Chapel Hill, Chapel Hill, NC 27599, USA; Department of Biochemistry and Molecular Biology, University of Arkansas for Medical Sciences, Little Rock, AR 72205, USA; Tsinghua-Peking Center for Life Sciences, Beijing Advanced Innovation Center for Structural Biology, and Beijing Frontier Research Center for Biological Structure, Tsinghua University, Beijing 100084, China; Department of Genetics, University of North Carolina at Chapel Hill School of Medicine, Chapel Hill, NC 27599, USA

**Author notes:** The authors wish it to be known that, in their opinion, the first two authors should be regarded as joint First Authors.

## Abstract

Trimethylation of histone H3 lysine 27 (H3K27me3) is important for gene silencing and imprinting, (epi)genome organization and organismal development. In a prevalent model, the functional readout of H3K27me3 in mammalian cells is achieved through the H3K27me3-recognizing chromodomain harbored within the chromobox (CBX) component of canonical Polycomb repressive complex 1 (cPRC1), which induces chromatin compaction and gene repression. Here, we report that binding of H3K27me3 by a Bromo Adjacent Homology (BAH) domain harbored within BAH domain-containing protein 1 (BAHD1) is required for overall BAHD1 targeting to chromatin and for optimal repression of the H3K27me3-demarcated genes in mammalian cells. Disruption of direct interaction between BAHD1^BAH^ and H3K27me3 by point mutagenesis leads to chromatin remodeling, notably, increased histone acetylation, at its Polycomb gene targets. Mice carrying an H3K27me3-interaction-defective mutation of Bahd1^BAH^ causes marked embryonic lethality, showing a requirement of this pathway for normal development. Altogether, this work demonstrates an H3K27me3-initiated signaling cascade that operates through a conserved BAH “reader” module within BAHD1 in mammals.

**Key Points:**

1. BAHD1^BAH^ is a functionally validated mammalian “reader” of H3K27me3, mediating BAHD1 targeting for gene silencing.
2. BAHD1^BAH^ connects H3K27me3 together with histone deacetylation, an integral step of gene silencing.
3. BAHD1^BAH^-mediated functional readout of H3K27me3 is essential for organismal development.

**Graphic abstract:** A mammalian H3K27me3-transduction pathway operates through an H3K27me3-specific ‘reader’ module (BAH) of BAHD1, which assembles a complex with corepressors (HDACs and others) for suppressing histone acetylation and repressing expression at Polycomb target genes.

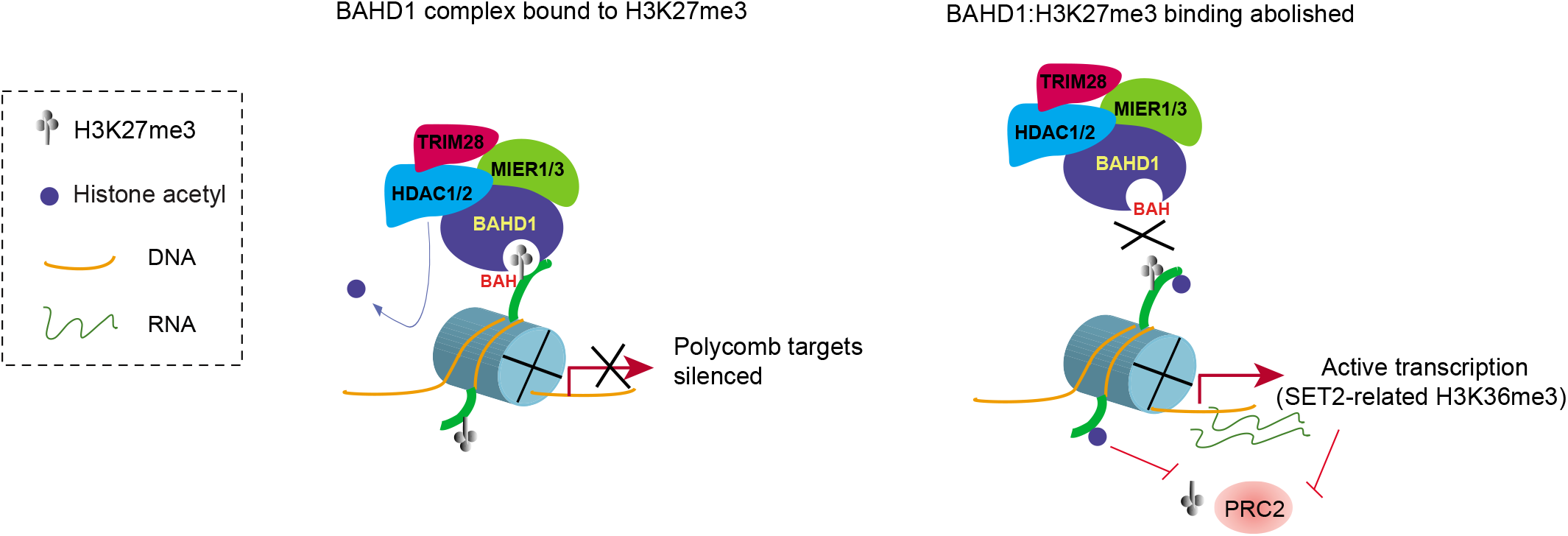

## INTRODUCTION

Trimethylation of histone H3 lysine 27 (H3K27me3), catalyzed by Polycomb repressive complex 2 (PRC2), is widely viewed to be crucial for regulation of gene silencing and imprinting, epigenomic and cellular states, cell fate determination, and embryogenesis (1–5). As a support, mutation of the histone H3K27 site or its modifying enzymes occur recurrently in a range of human diseases, such as cancer and developmental syndrome (6–9). A prevalent model for functional transduction of H3K27me3 in mammals is that H3K27me3 is read and bound by chromodomain harbored within the chromobox (CBX) component of canonical Polycomb repressive complex 1 (cPRC1), which can induce gene silencing at least partly through mono-ubiquitination of histone H2A lysine 119 (H2AK119ub1) (2,3,10,11) and/or a phase separation-mediated chromatin compaction mechanism (12–14). However, emerging evidence has shown that, in mammalian cells, H3K27me3 is also ‘sensed’ and bound by a second class of “reader” modules termed Bromo-Adjacent Homology (BAH), which exist within chromatin regulators such as <u>BAH</u> <u>D</u>omain Containing <u>1</u> (BAHD1) (15,16) and <u>BAH</u> and <u>C</u>oiled-<u>c</u>oil Domain-containing Protein <u>1</u> (BAHCC1) (17). In particular, it was recently demonstrated that disrupting the direct binding of the BAHCC1 BAH motif (BAHCC1^BAH^) to H3K27me3 via point mutagenesis leads to significant de-repression of the H3K27me3-demacated genes (17). Mechanistically, BAHCC1 interacts with histone deacetylases (HDACs), thereby linking H3K27me3 with histone deacetylation, an integral step of gene repression (17). Interestingly, BAHD1 also interacts with HDACs and mediates heterochromatin formation in mammalian cells (15,18). Although the BAH domain within BAHD1 (BAHD1^BAH^) was previously shown to bind H3K27me3 directly (16), the requirement of such an interaction for target gene regulation yet remains uncharacterized to date; a multifunctional scaffold role of BAHD1 was previously proposed for inducing gene repression, which involves the H3K9me3-specific readers and other BAHD1-associated silencing factors (15,18). It shall be noted that, in plant and fungus, the cPRC1 pathway gene(s) is absent and, instead, a set of the plant/fungus-specific BAH-containing proteins were recently identified, which directly engage H3K27me3 for mediating Polycomb gene silencing (19–24). These observations highlight biological significance and cross-species conservation of the BAH module-based functional readout of H3K27me3. Dissecting the molecular underpinning of chromatin modification-initiated signal transduction shall help understand the fundamentals of (epi)genomic and transcriptomic regulation.

Using independent mammalian model systems such as 293 cells, primary mouse embryonic fibroblast (MEF) cells and the genetically modified knockin mice, we here aim to determine the role of the H3K27me3-binding BAHD1^BAH^ in target gene regulation. Our findings, based on mutagenesis of BAHD1^BAH^, substantiate a critical requirement of BAHD1^BAH^:H3K27me3 interaction for overall BAHD1 targeting to chromatin and for maintaining local histone deacetylation, resulting in optimal repression of the BAHD1-bound Polycomb target genes. Importantly, the readout of H3K27me3 by BAHD1^BAH^ is essential for embryogenesis and survival in mice. We thus favor a view that the functional transduction of H3K27me3 via BAH-directed molecular pathways is evolutionarily conserved across the domain of Eukaryota, including animal, which operate to ensure appropriate epigenomic and transcriptomic patterns during organismal development.

## MATERIALS AND METHODS

### Plasmids

Full-length cDNA of human BAHD1 (a kind gift of H. Bierne, L’Institut Micalis, France), together with a 3×FLAG tag fused to its C-terminal, was cloned into the XhoI and NotI site of the pPyCAGIP vector (carrying IRES-Puro; a kind gift of I. Chambers). Point mutation was generated by site-directed mutagenesis. Information of primers used in the work is listed in Supplementary Table 6. Correctness of all plasmids was verified by Sanger sequencing before use.

### Cell lines and tissue culture

HEK293 cells (American Tissue Culture Collection, #CRL-1573) were maintained under the vendor-recommended culture condition. Cells were transfected by Lipofectamine 2000 (Invitrogen) with the pPyCAGIP empty vector or that carrying wild-type or mutant 3×FLAG-BAHD1. Two days post-transfection, HEK293 cells were subjected to drug selection in the medium with 2 μg/ml puromicin (Invitrogen) for 14 days. The stable expression lines were continuously maintained in the medium with puromycin. Authentication of cell line identity, including parental and their derived lines, was ensured by the Tissue Culture Facility (TCF) affiliated to UNC Lineberger Comprehensive Cancer Center with the genetic signature profiling and fingerprinting analysis (25). Each month, a routine examination of cells for mycoplasma contamination was conducted by using the detection kit (Lonza). Cells with less than 10 times of passages were used in the study.

### RNAi for gene knockdown (KD)

Cells were transfected with siRNA using the Lipofectamine RNAiMAX reagent according to manufacturer’s instruction. The Silencer™ Select Pre-Designed siRNA for human BAHD1 (Thermo Fisher 4392422, siRNA # s22604) was ordered and used per vendor’s guideline.

### Quantitative RT-PCR (RT-qPCR)

RT-qPCR was carried out as previously described (17,26), with the primer sequence listed in Supplementary Table 6.

### Co-immunoprecipitation (CoIP) and peptide pulldown

Whole cell lysates were prepared by using the NP40 lysis buffer (50mM Tris-HCl pH 8.0, 150mM NaCl, 1% IGEPAL® CA-630, and 1× protease inhibitor cocktail added freshly), followed by brief sonication and high-speed centrifugation at 4°C. For CoIP, cell lysates were incubated with antibodies on a rotator overnight in a cold room, followed by addition of protein A/G beads for additional 2 hrs. Beads were washed three times in NP40 lysis buffer, resuspended in 50 μl of 1× SDS sample buffer, and boiled at 90 degree for 5 min before loading onto SDS-PAGE gel. We conducted the biotinylated peptide pulldown using total cell lysate according to a previously described protocol (17,26).

### Antibodies and immunoblotting

The information of antibodies used for immunoblotting, immunofluorescence, CoIP and ChIP, ChIP-seq and CUT&RUN can be found in Supplementary Table 6. Protein samples, either total cell lysates or samples from pulldown or immunoprecipitation, were loaded onto SDS-PAGE gels for immunoblotting as described before (17,26). Western blotting was performed with standard protocols using the PVDF membrane and signals visualized with ImageQuant LAS 4000 Luminescent Image Analyzer (GE Healthcare).

### Immunofluorescence

Cells were fixed in 4% paraformaldehyde for 15 min at room temperature, followed by incubation in 1× PBS containing 0.25% of Triton X-100 for 10 min to permeabilize the cells. Fixed cells were co-stained with primary antibodies and then with Alexa-488/594 conjugated secondary antibodies (Life Technologies) as described before (17,26). Nuclei were stained with 4, 6-diamino-2-phenylindole (DAPI, 0.1 μg/ ml). Fluorescent signals were detected and images were taken by an FV1000 confocal microscope (Olympus, UNC Imaging Core).

### Chromatin fractionation assay

Two million of cells were collected and re-suspended in 100 μl of ice-cold CSK buffer (10 mM PIPES, pH 7.0; 0.1% Triton X-100; 300 mM sucrose; 100 mM NaCl; 3 mM MgCl_2_), followed by incubation on ice for 20 minutes. Samples were then centrifuged at 1,300g for 5 minutes at 4°C. The supernatant was collected as the soluble fraction and the pellet as the chromatin-associated fraction.

### Compound

The catalytic inhibitor of EZH2 and EZH1, UNC1999, was dissolved in dimethylsulfoxide (DMSO) as 5 mM stock solution and used as previously described (27,28). The histone deacetylase (HDAC) inhibitor, CI-994, was used as we described before (17). The cell treatment condition for UNC1999 and CI-994 in this study was 5 μM as the final concentration for three and five days, respectively.

### RNA-seq and Data Analysis

Total RNA was extracted using the RNeasy Mini Kit (QIAGEN) and RNA-seq libraries were prepared using the TruSeq RNA Sample Preparation Kit v2 (Illumina) as previously described (17,29). Multiplexed RNA-seq libraries were pooled and subjected for deep sequencing on an Illumina Hi-Seq 2500 or 4000 platform (UNC High-throughput Sequencing Facility). For data analysis, RNA-seq reads were first mapped using MapSplice (30) and quantified using RSEM (31). Read counts were then upper-quantile normalized and log2 transformed. Raw read counts were used for differential gene expression analysis by DESeq (32).

### Gene Ontology (GO) analysis and Gene Set Enrichment Analysis (GSEA)

GO and GSEA analyses were conducted as previously described (27,29,33). GSEA was carried out by using the GSEA_4.0.3 software (34) and the Molecular Signatures Database v6.2 (MSigDB, http://www.broadinstitute.org) for testing putative enrichment of hallmark gene sets (H), curated gene sets (C2) and oncogenic gene sets (C6).

### Chromatin immunoprecipitation (ChIP) followed by qPCR (ChIP-qPCR)

ChIP-qPCR was carried out as previously described (35,36). In brief, one million of cells were harvested and cross-linked with 1% of formaldehyde at room temperature for 10 minutes, followed by adding 2.5 M of glycine to quench crosslinking. Cells were sonicated by Bioruptor (Diagenode) and the cleared chromatin samples incubated with antibody-conjugated Dynabeads (Invitrogen) at 4 degree for overnight. Beads bound with chromatin were subject to extensive washing, followed by elution, de-crosslink overnight at 65 degree, and sequential digestion first by proteinase K (Thermo Fisher Scientific) and then RNase A (Roche). The ChIP DNA was then purified with MinElute PCR Purification Kit (QIAGEN). The samples of ChIP DNA and matched input DNA (similarly processed) were diluted in water and used for ChIP-qPCR. ChIP signals, generated from three independent experiments, were normalized to input signals and presented as mean ± SD. Student’s t-test was used for data analysis. The sequence of primers used for ChIP-qPCR is available in supplementary table.

### ChIP followed by sequencing (ChIP-seq) and data analysis

ChIP-seq was performed as previously described (27,29,33), with a multi-step procedure that involves the cell fixation and sonication, antibody incubation, washing, ChIP DNA recovery, library preparation, and deep sequencing (UNC High-throughput Sequencing Facility). The standard data analysis of ChIP-Seq data alignment, filtration, peak calling and assignment, and cross-sample comparison was performed as described before (27,29,33) with slight modification. In brief, ChIP-seq reads were aligned to human genome build GRCh37/hg19 using STAR version 2.7.1a (37). Alignment files in the bam format were transformed into read coverage files (bigwig format) using DeepTools (38). Genomic binding profiles were generated using the deepTools “bamCompare” functions with options [--operation ratio ---pseudocount 1 −binSize 10 --extendReads 250] and normalized to the matched input profile. The resulting bigWig files were visualized in the Integrative Genome Viewer (IGV). Heatmaps were generated using the deepTools “computeMatrix” and “plotHeatmap” functions. The genomic regions covering from the transcription start site (TSS) to transcription end site (TES) of all genes annotated in the human genome (NCBI Reference Sequences [RefSeq]; GRCh37/hg19) were obtained from UCSC table browser and used for heatmap and averaged signal plotting.

### CUT&RUN followed by sequencing

CUT&RUN (39) was performed with a commercially available kit according to manufacturer’s instruction (EpiCypher CUTANA™ pAG-MNase for ChIC/CUT&RUN, Cat# 15-1116). Briefly, one million of cells were first collected, washed in the CUT&RUN wash buffer (20 mM HEPES, pH 7.5, 150 mM NaCl, 0.5 mM spermidine, 1× Complete Protease Inhibitor Cocktail), and then bound to the activated ConA beads (Bangs Laboratories BP531). Next, the cell:bead sample was incubated with antibodies against the protein target (1:100 dilution) and then permeabilized in the digitonin-containing buffer (CUT&RUN wash buffer plus 0.01% digitonin), which was then followed by washing in the digitonin buffer, incubation with pAG-MNase, and another washing in the digitonin buffer to remove the unbound pAG-MNase. Following the final wash, cells were subjected to digestion following the pAG-MNase activation by addition of the pAG-MNase digestion buffer (digitonin buffer plus 2mM CaCl2), followed by incubation on nutator for 2 hours at 4°C. Solubilized chromatin was then released using the CUT&RUN stop buffer (340 mM NaCl, 20 mM EDTA, 4 mM EGTA, 50 μg/ml RNase A, 50 μg/ml glycogen) and DNA purification was carried out with the PCR cleanup kit (NEB, Monarch PCR & DNA Cleanup Kit, T1030). About 10ng of the purified CUT&RUN DNA was used for preparation of multiplexed libraries with the NEB Ultra II DNA Library Prep Kit per manufacturer’s instruction (NEB #E7103). Sequencing was conducted using an Illumina NextSeq 500 Sequencing System (available from the core facility of UNC Pharmacology Department).

### CUT&RUN data analysis

Fastq files were mapped to the reference genome (hg19) using bowtie2.3.5 (40). The non-primary alignment, PCR duplicates, or blacklist regions were removed from aligned data by Samtools (v1.9), Picard “MarkDuplicates” function (ver 2.20.4), and bedtools (v2.28.0), respectively. Peak calling was performed using MACS2 (macs2 callpeak −f BAMPE −g hs/mm --keep-dup 1) (41). Distribution of peaks was calculated by “annotatePeaks.pl” function of HOMER (Hypergeometric Optimization of Motif Enrichment) (42). Deeptools (v3.3.0) was used to generate bigwig files, heatmaps, and averaged plotting of CUT&RUN signals (38). Genomic binding profiles were generated using the deepTools “bamCompare” functions.

### Mass spectrometry–based analysis of BAHD1-containing protein complexes

As previously described (17,26), nuclear extract was prepared using the Dignam protocol from the nuclei of HEK293 cells that stably express 3xFLAG-tagged BAHD1, with parental HEK293 cells as a negative control. Following dialysis against low-salt buffer (150 mM NaCl, 20mM Hepes, pH 7.90, 0.025% IGEPAL® CA-630, 0.2 mM EDTA, 10% glycerol, 1 mM DTT, 0.2 mM PMSF), nuclear extracts were centrifuged at high speed to remove precipitation and supernatants incubated with FLAG M2 beads (Sigma). After extensive washing, the resin and bound proteins from both experimental and negative control cells were subjected to mass spectrometry-based protein identification (carried out at Proteomics Core facility, University of Arkansas for Medical Sciences).

### Generation of the *Bahd1^W659G^* knock-in allele by CRISPR/cas9-mediated gene editing in the C57BL/6J mouse strain

Guide RNAs, which target the site close to the *Bahd1* W659 codon in exon 5, were identified using Benchling software. Three guide RNAs were selected for initial activity testing. Here, the selected guide RNAs were cloned into a T7 promoter vector followed by in vitro transcription and spin column purification. Functional testing was performed by cotransfecting a mouse embryonic fibroblast cell line with guide RNA and Cas9 protein. Cas9 protein was expressed and purified by the UNC Protein Expression Core Facility. The guide RNA target site was amplified from transfected cells and analyzed by T7 Endonuclease I digestion of re-annealed PCR products (New England Biolabs). The guide RNA eventually selected for genome editing in embryos was Bahd1-sg47B (protospacer sequence 5’-GCCTGTAATACCACAGG −3’). A donor oligonucleotide (Bahd1-W659G-T), with the sequence of 5’-CTAGCCATGACCTCTTCTCTTGTCCCCTAGGAGAGCTGATGATGAGCCTCCTG**<u>GGC</u>**TATTACAGGCCTGAGCACTTACAGGGAGGTCGCAGCCCCAGCATGCATGAGG-3’, was designed for homologous recombination to insert the TGG to GCC (bold/underlined in oligo sequence) mutation of Tryptophan 659 to Glycine.

C57BL/6J zygotes were microinjected with 400 nM Cas9 protein, 20 ng/ul guide RNA and 50 ng/ul donor oligonucleotide. In vitro transcribed, purified guide RNA was diluted in microinjection buffer (5 mM Tris-HCl pH 7.5, 0.1 mM EDTA), heated at 95 degree for 3 min, and placed on ice prior to addition of Cas9 protein. The mixture was then incubated at 37 degree for 5 min and placed on ice, after which the donor oligonucleotide was added. The final mixture was filtered through a 0.2 uM filter and held on ice prior to microinjection.

All animal experiments were approved by and performed in accord with the guideline of Institutional Animal Care and Use Committee at the University of North Carolina (UNC) at Chapel Hill, USA. Microinjected embryos were implanted in recipient pseudopregnant females. Resulting pups were screened by PCR and sequencing for the presence of the mutant allele. Of 17 pups born that survived to genotyping, 5 showed clear evidence of the W659G mutant allele. Independent male founders were mated to wild-type C57BL/6J females, and transmitted the mutant allele through the germline to offspring.

### Generation of primary mouse embryonic fibroblast (MEF) cells

The *Bahd1^W659G^* heterozygous mice were set up for breeding and the E13.5 embryos harvested. The isolation and establishment of primary MEF cells were conducted as previously described (43). In brief, unwanted organs of embryos were removed, followed by tissue homogenization and enzymatic dissociation of cells. Genotypes of cells were determined by PCR and direct sequencing. Cells were then cultured in the DMEM base medium supplemented with 15% of FBS for 25 passages to establish stable MEF cell lines.

### Statistical analysis

Experimental data are presented as mean ± SD of three independent experiments. Statistical analysis was performed with Student’s t test for comparing two sets of data with assumed normal distribution. *P* < 0.05 was considered significant. Statistical significance levels are denoted as follows: \**P*<0.05; \*\**P*<0.01; \*\*\**P*<0.001; \*\*\*\**P*<0.0001. No statistical methods were used to predetermine sample size.

## RESULTS

### Specific interaction between the BAH domain of BAHD1 (BAHD1^BAH^) and H3K27me3 is required for overall binding of BAHD1 to chromatin

A conserved BAH module located at the C-terminus of BAHD1 (BAHD1^BAH^; Supplementary Figure 1A) was shown to bind H3K27me3 directly (16), an interaction that we confirmed by immunofluorescence (IF) of H3K27me3 and 3xFLAG-tagged BAHD1 (f-BAHD1) in the HEK293 stable expression cells (Supplementary Figure 1B-C). When compared to BAHCC1^BAH^, another H3K27me3-specific “reader” module (17), BAHD1^BAH^ displays considerable overall sequence divergence but retains conservation at so-called H3K27me3-”caging” residues (see stars in Supplementary Figure 1A). By CoIP, we further showed that BAHD1 associates with cellular H3K27me3 and not the other examined histone trimethylation, including H3K4me3, H3K9me3, H3K36me3, H3K79me3 and H4K20me3 (Figure 1A). To assess the requirement of such a BAHD1^BAH^:H3K27me3 interaction for tethering BAHD1 to chromatin, we established the HEK293 stable cells that expressed the comparable level of BAHD1, either wildtype (WT) or with point mutation of a conserved “caging” residue within BAHD1^BAH^ (W667G or Y669A, see also stars in Supplementary Figure 1A; Figure 1B, top). Relative to WT, either “caging” mutation of BAHD1^BAH^ disrupted the H3K27me3 binding in the histone peptide pulldown assay (Figure 1B, bottom). Additionally, immuoblotting with the cell fractionation samples detected a significant decrease in the chromatin association by BAHD1 and a simultaneous increase of BAHD1 in the soluble fraction (Figure 1C). Thus, direct engagement of H3K27me3 by BAHD1^BAH^ is required for overall chromatin targeting of BAHD1 in cells.

**Figure 1.**
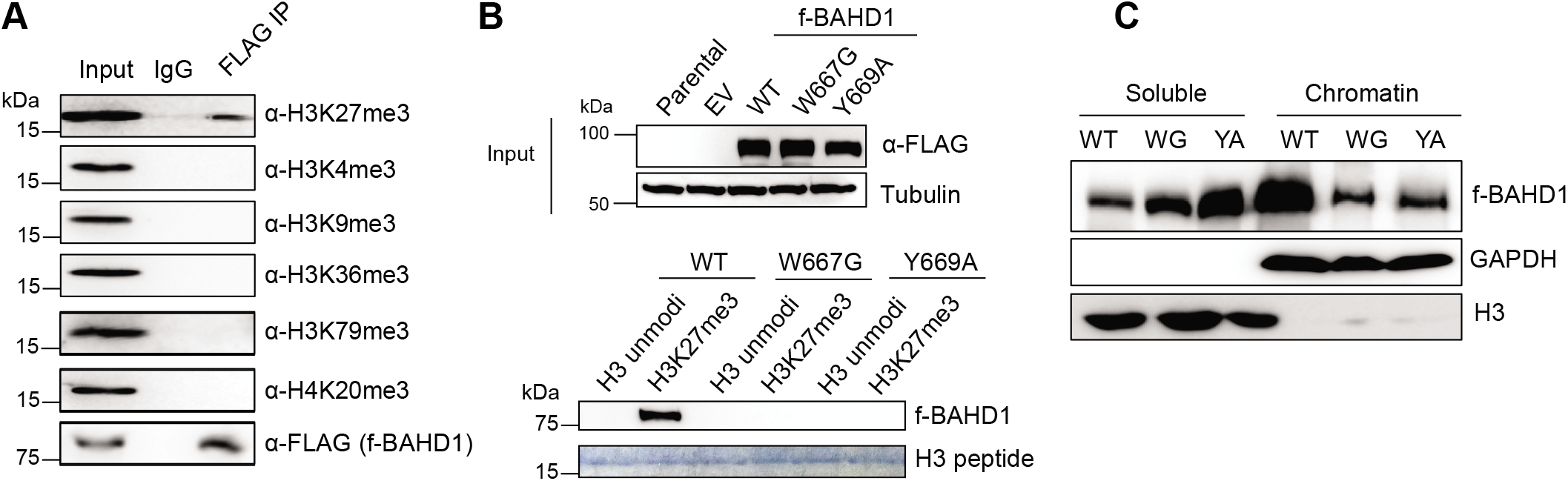
H3K27me3 binding by the BAHD1 BAH module (BAHD1^BAH^) contributes significantly to overall chromatin association of BAHD1. **(A)** Co-IP between FLAG-tagged BAHD1 (f-BAHD1) and the indicated histone methylation in HEK293 cells. (**B**) Top: anti-FLAG immunoblotting using total cell extracts of HEK293 cells, either parental or those with stable expression of empty vector (EV) or f-BAHD1, WT or BAH-mutated (W667G or Y669A). Bottom: Pulldown using the biotinylated histone H3 peptide, either unmodified or harboring H3K27me3, and total cell extracts that contain WT or BAH-mutated f-BAHD1, followed by anti-FLAG immunoblot. (**C**) Immunoblotting for f-BAHD1 using the chromatin-bound (right) and soluble (left) fractions prepared from the HEK293 cells expressing either WT or BAH-mutated (W667G or Y669A) f-BAHD1. GAPDH and H3 served as the control of fractionation.

### BAHD1^BAH^ mediates BAHD1 targeting to the H3K27me3-marked Polycomb targets

Next, we performed H3K27me3 ChIP-seq and mapped the genome-wide binding of f-BAHD1 with CUT&RUN in HEK293 cells. The called peaks of f-BAHD1 are mostly located at promoter, intergenic and intron regions (Figure 2A) whereas a negative control experiment of CUT&RUN by using parental HEK293 cells without f-BAHD1 expression did not produce the enriched signals (Supplementary Figure 2A-B). There is a significant overlap between overall bindings of BAHD1 and H3K27me3 (Figure 2B-C). Heatmaps showed that genes with the more BAHD1 binding also tend to carry more H3K27me3 (Figure 2C), as exemplified by what was observed at the classic Polycomb target genes such as IGF2, FOXB1 and CCND2 (Figure 2D, the top and bottom tracks). Compared to WT, overall genome binding of the H3K27me3-binding-defective mutant, either BAHD1^W667G^ or BAHD1^Y669A^, was found significantly reduced (Figure 2E and the two middle tracks of Figure 2D), in agreement with the decreased chromatin association of these BAH mutants (Figure 1C). These results lend strong support for a critical involvement of BAHD1^BAH^ in targeting BAHD1 to the H3K27me3-associated genomic loci.

**Figure 2.**
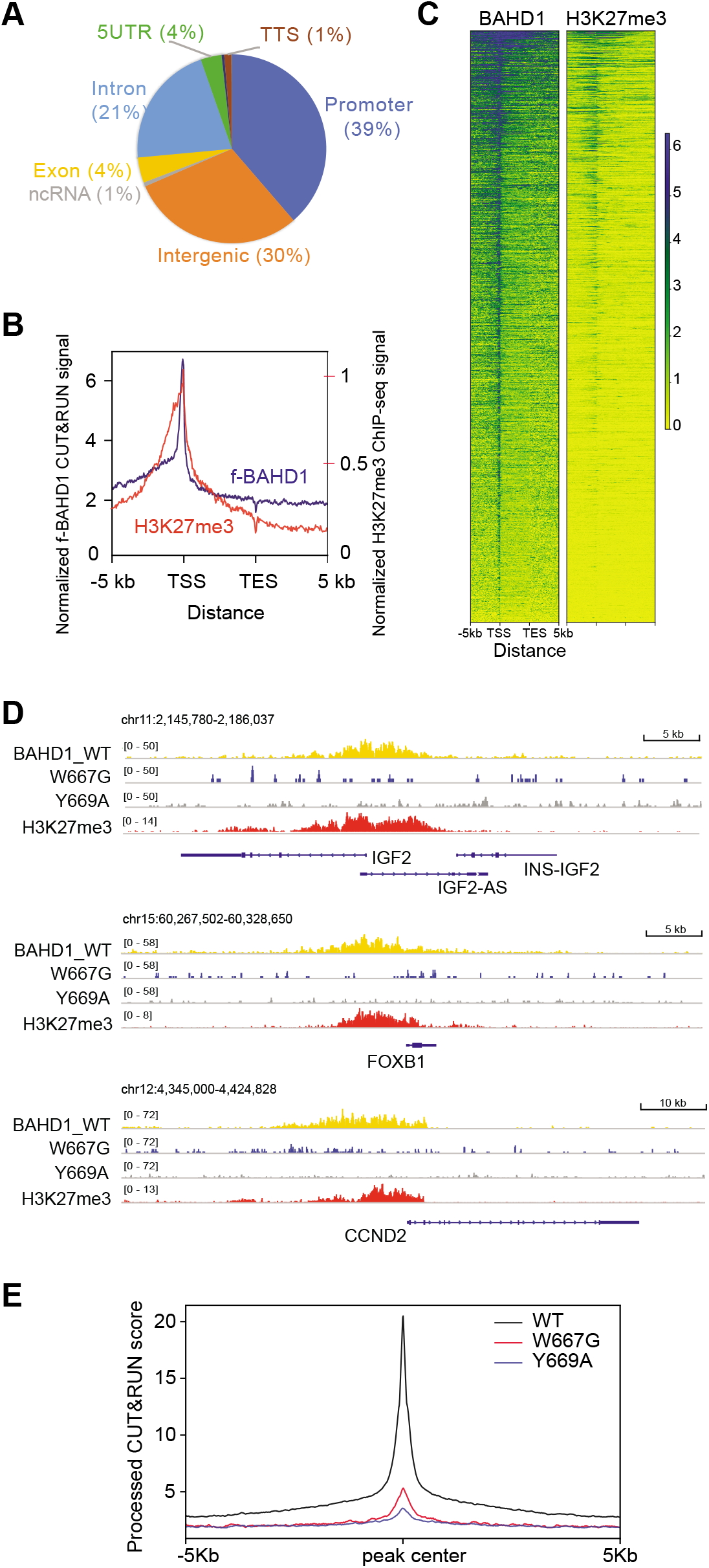
BAHD1^BAH^-mediated H3K27me3 engagement is required for BAHD1’s recruitment to Polycomb target genes. **(A)** Pie chart showing distribution of the called f-BAHD1 CUT&RUN peaks among the indicated genomic features in HEK293 cells. **(B-C)** Plotting of averaged intensity **(B)** and heatmaps **(C)** of f-BAHD1 CUT&RUN and H3K27me3 ChIP-seq signals across ±5 kb from transcription start site (TSS) and transcription end site (TES) of target genes in HEK293 cells. (**D**) IGV views showing the called peaks of WT or BAH-mutated (W667G or Y669A) f-BAHD1, as well as H3K27me3, at the indicated gene in HEK293 cells. **(E)** Averaged CUT&RUN signals (y-axis) of the indicated WT and mutant f-BAHD1 across ±5 kb from the peak center (x-axis) in HEK293 stable expression cells.

### BAHD1^BAH^ is critical for the BAHD1-mediated Polycomb gene repression in HEK293 cells

We next sought to determine the requirement of an H3K27me3-binding BAHD1^BAH^ motif for regulation of target gene expression. Towards this end, we first identified siRNA that allowed an almost complete depletion of BAHD1 in HEK293 cells (Figure 3A; Figure 3B, lane 2 vs. 1). To assess the role of BAHD1^BAH^ per se, we next generated a BAHD1 cDNA for rescuing the endogenous BAHD1 knockdown (KD) due to its resistance to the used BAHD1-targeting siRNA (Figure 3C), as demonstrated by immunoblotting post-transfection of siRNA into cells with stable expression of such exogenous f-BAHD1, either WT or an H3K27me3-binding-defective mutant (Figure 3B, lanes 4, 6 and 8 versus lane 2). Utilizing such a validated rescue system, we subsequently performed RNA-seq profiling, which revealed significant transcriptomic alterations caused by the BAH mutation (W667G; Figure 3D and Supplementary Table 1). Relative to WT, BAHD1^W667G^ predominantly caused gene expression upregulation (Figure 3D, red). Both Gene Set Enrichment Analysis (GSEA; Figure 3E) and Platform for Integrative Analysis of Omics data (PIANO (44); Supplementary Figure 3A) further uncovered that, relative to WT, BAHD1^W667G^ is most significantly associated with de-repression of genesets previously known to be marked by H3K27me3, repressed by Polycomb factor (PRC2 or PRC1), or related to organismal development (Figure 3E-F and Supplementary Figure 3A-B). By RT-qPCR, we verified the marked reactivation of the tested Polycomb targets, which included a classic imprinted gene (IGF2), development genes (SOX8, WNT7B and MEIS3) and IGFBP2 (H3K27me3-marked), post-KD of endogenous BAHD1 relative to mock (Figure 3G, red vs. black); such a change was found to be reversed due to pre-rescue with exogenous WT BAHD1, but not either one of the H3K27me3-binding-defective mutants (BAHD1^W667G^ or BAHD1^Y669A^; Figure 3G, cyan and blue vs. yellow and black). Meanwhile, neither depletion of BAHD1 nor expression of its mutant altered the protein level of Polycomb proteins or global H3K27me3 (Figure 3H and Supplementary Figure 3C). Based on genomic profiling data (CUT&RUN and RNA-seq), we have also defined those BAHD1/H3K27me3 co-bound target genes that exhibit a repression pattern in a BAHD1^BAH^-dependent manner (Figure 3I and Supplementary Table 2), and this gene signature includes a number of classic Polycomb gene targets such as lineage-specific transcription factors and developmental regulators. Altogether, these observations highlight a role of direct BAHD1^BAH^:H3K27me3 interaction in the BAHD1-related Polycomb gene silencing.

**Figure 3.**
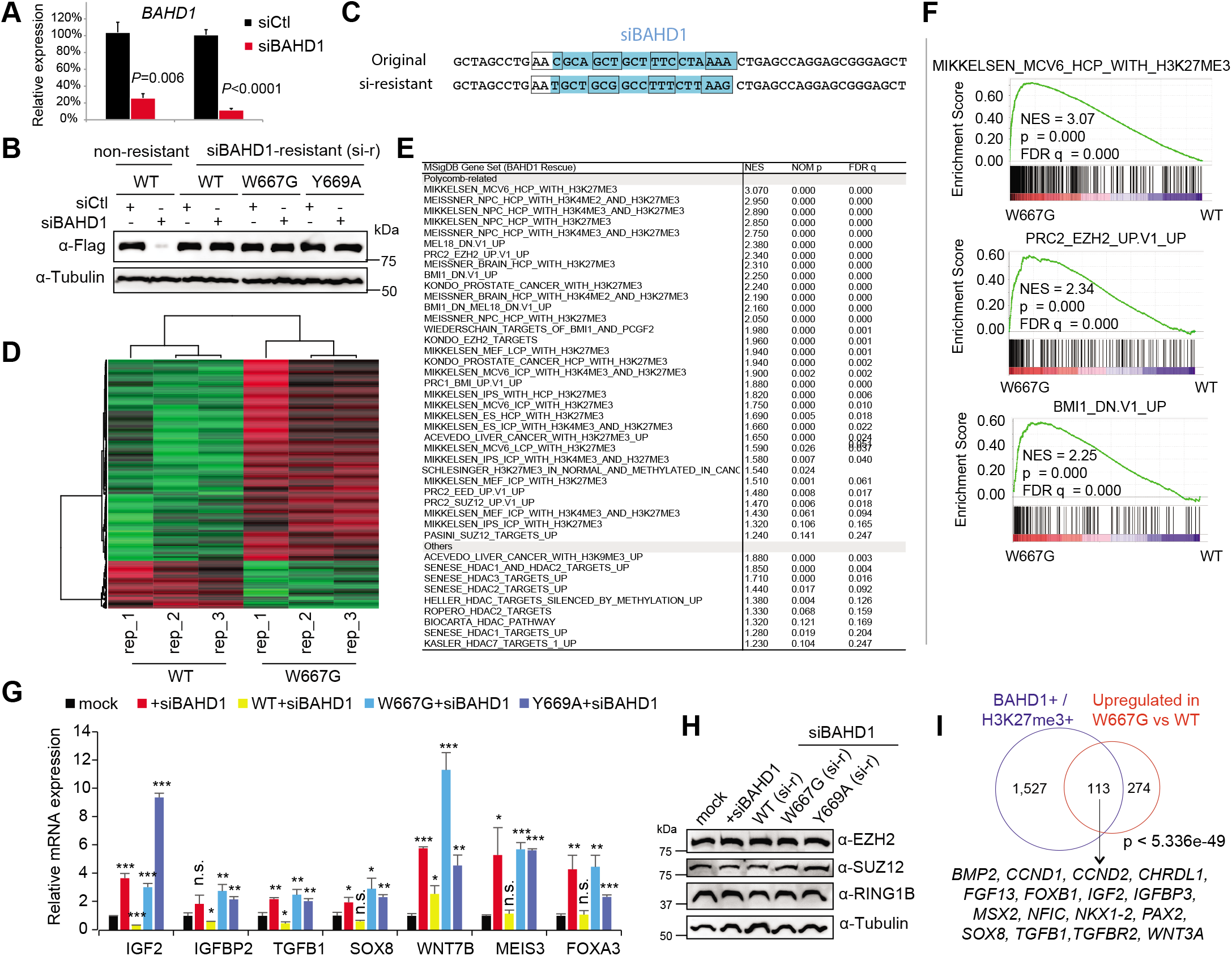
Transcriptomic profiling demonstrates an essential involvement of BAHD1^BAH^ in BAHD1-induced gene repression in HEK293 cells. **(A)** RT-qPCR of endogenous *BAHD1* after siRNA-mediated knockdown (KD) in HEK293 cells (n = 3 biologically independent samples). Two different sets of BAHD1 primers were used. Data are presented as mean ± SD. (**B**) Immunoblotting of exogenous f-BAHD1, either WT or with a mutation of BAHD1^BAH^ (W667G or Y669A), post-transduction of the BAHD1-targeting siRNA (siBAHD1) or scramble control (siCtl). Left and right panels showed the HEK293 cell lines with stable expression of f-BAHD1 that was efficiently depleted by siBAHD1 (left) or designed to be siBAHD1-resistant (“si-r”; right, see also panel C). (**C**) The BAHD1 cDNA sequence targeted by the used BAHD1-targeting siRNA (siBAHD1 as shown in blue on the top; see also lanes 2 vs 1 in panel B), and a designed siBAHD1-resistant BAHD1 form that harbors seven silent point mutations of the siBAHD1-targeting site (bottom; see also lanes 3-8 of panel B). Each box indicates a codon. (**D**) Heatmap showing the relative expression of differentially expressed genes (DEGs) identified in HEK293 cells with rescue of WT BAHD1, compared to its W667G mutant, followed by endogenous BAHD1 KD by siBAHD1 (n=3 biological replicates per group). The thresholds of DEGs are adjusted *P* value (adj. p) < 0.01, fold-change (FC) >2.0 and transcript baseMean value > 10 based on RNA-seq. Color bar, log2(FC). (**E**) Summary of GSEA using RNA-seq profiles of HEK293 cells that expressed an H3K27me3-binding-defective mutant of BAHD1 (W667G), compared to WT. NES, normalized enrichment score. (**F**) GSEA revealing that, relative to WT, the BAH mutation (W667G) of BAHD1 is positively correlated to reactivation of the indicated Polycomb gene signature associated with H3K27me3, PRC2 (EZH2) or PRC1 (BMI1). (**G**) RT-qPCR of the indicated BAHD1-repressed gene 48 hours post-transduction of the scramble control (mock) or BAHD1-targeting siRNA (siBAHD1) into parental HEK293 cells (black and red), as well as post-transduction of siBAHD1 into HEK293 cell lines pre-rescued with exogenous siBAHD1-resistant BAHD1, either WT or BAH-mutated (W667G or Y669A). Data of three independent experiments are presented as mean ± SD after normalization to GAPDH and then to mock. n.s., not significant; * *P* < 0.05; ** *P* < 0.01; *** *P* < 0.001. (**H**) Immunoblotting for Polycomb proteins (EZH2, SUZ12 and RING1B) in the indicated HEK293 cells used in the panel **G**. (**I**) Venn diagram showing overlap between the BAHD1/H3K27me3 co-bound genes (left) and DEGs derepressed due to the BAHD1 W667G mutation, relative to WT (right), in HEK293 cells. Below lists the Polycomb target examples.

### H3K27me3 and BAHD1 co-repress a set of Polycomb target genes in HEK293 cells

To further define genes repressed by H3K27me3 per se, we employed a catalytic inhibitor of PRC2, UNC1999 (27,28), to ‘erase’ global H3K27me3 in HEK293 cells (Figure 4A), followed by RNA-seq (Supplementary Table 3). UNC1999 mainly induced gene derepression (Figure 4B). As expected, these UNC1999-activated transcripts were most enriched with those previously known to be the H3K27me3-demarcated, Polycomb-targeted, and/or development-associated ones (Supplementary Figure 4A-B). Repressive effect by PRC2-catalyzed H3K27me3 was further verified by RT-qPCR of several Polycomb targets post-treatment of UNC1999 (Figure 4C). Importantly, differentially expressed genes (DEGs) reactivated by UNC1999 significantly overlapped with those derepressed due to the H3K27me3-interaction-disrupting mutation of BAHD1^BAH^, relative to WT (Figure 4D). In consistence, UNC1999 treatment also significantly dissociated BAHD1 off the chromatin (Figure 4E). These results thus lent a strong support for an axis of H3K27me3→BAHD1^BAH^ →repression.

**Figure 4.**
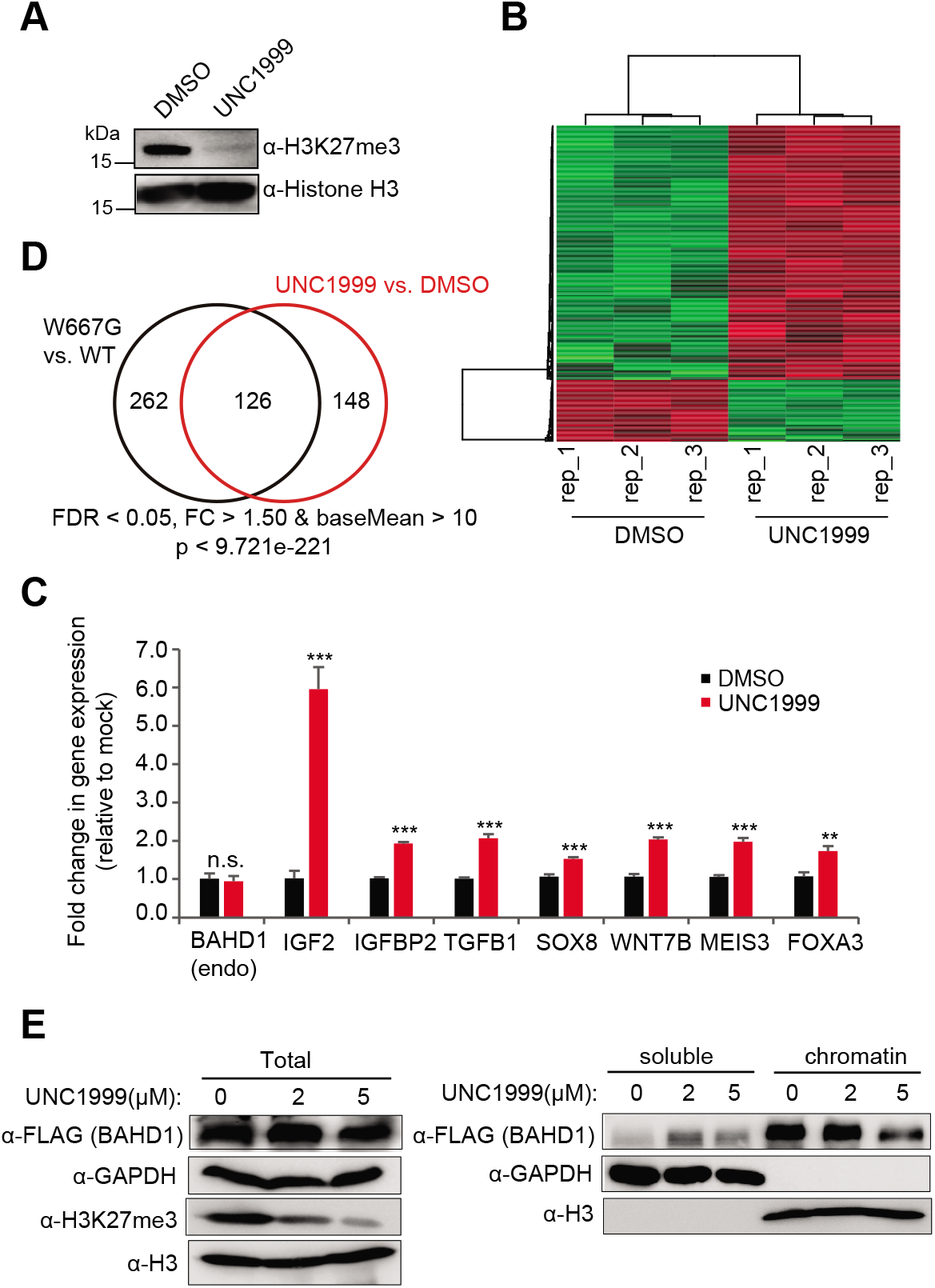
H3K27me3 and BAHD1 co-repress Polycomb target genes. (**A**) H3K27me3 and total H3 immunoblots using HEK293 cells post-treatment with 5 uM of DMSO or UNC1999 for three days. (**B**) Heatmap showing the relative expression of DEGs in HEK293 cells treated with 5 uM of UNC1999 (right) relative to DMSO (left) for three days, as revealed by RNA-seq (n=3 biological replicates per group). (**C**) RT-qPCR of BAHD1 and the indicated BAHD1-repressed target gene in HEK293 cells post-treatment with 5 uM of UNC1999 versus DMSO for three days (n = 3 biologically independent samples). Data are presented as mean ± SD after normalization to GAPDH and then to mock-treated. n.s., not significant; ** *P* < 0.01; *** *P* < 0.001. **(D)** Venn diagram showing significant overlap between the DEGs upregulated due to the W667G-mutated BAHD1 relative to WT (left) and those upregulated due to UNC1999 treatment relative to DMSO (right) in HEK293 cells. (**E**) Immunoblotting for f-BAHD1 using the total extract (left) or the chromatin-bound and soluble fractions (right) of the HEK293 cells post-treatment of UNC1999, relative to DMSO. GAPDH and H3 served as the control of fractionation.

### Disruption of the BAHD1^BAH^:H3K27me3 interaction results in chromatin remodeling, notably the histone acetylation increase, at the BAHD1/H3K27me3 co-targeted genes

We also performed f-BAHD1 pulldown in the HEK293 stable expression cells and the subsequent mass spectrometry-based analysis revealed a complex that comprises HDAC1, HDAC2 and MIER1/3 (Figure 5A and Supplementary Table 4), consistent with a previous report (18). Both CoIP and IF demonstrated a strong association of BAHD1 with HDACs (Figure 5B-C). Additionally, the aromatic “cage” mutations of BAHD1^BAH^ that disrupted direct H3K27me3 binding by BAHD1 did not affect the BAHD1:HDAC interaction (Figure 5D, WT versus WG or YA), indicative of different protein interfaces involved in the two interaction events. Notably, GSEA analyses of RNA-seq profiles also showed a positive correlation of HDAC target gene reactivation with BAHD1^W667G^ relative to WT (Figure 5E; Supplementary Figure 3A), as well as UNC1999 treatment relative to mock (Supplementary Figure 4A). By ChIP-qPCR, we detected the HDAC2 binding at the examined BAHD1 target genes (Figure 5F, red vs black). Such binding of HDAC2 to BAHD1 targets was found to be significantly reduced in HEK293 cells expressing the BAH mutant of BAHD1 (W667G or Y669A), compared to WT (Figure 5F, blue/yellow vs red). Treatment of HEK293 cells with the HDAC inhibitor CI-994 led to significant reactivation of BAHD1 target genes (Figure 5G). These observations highlight an important role for the BAHD1:HDAC complex in mediating Polycomb gene silencing.

**Figure 5.**
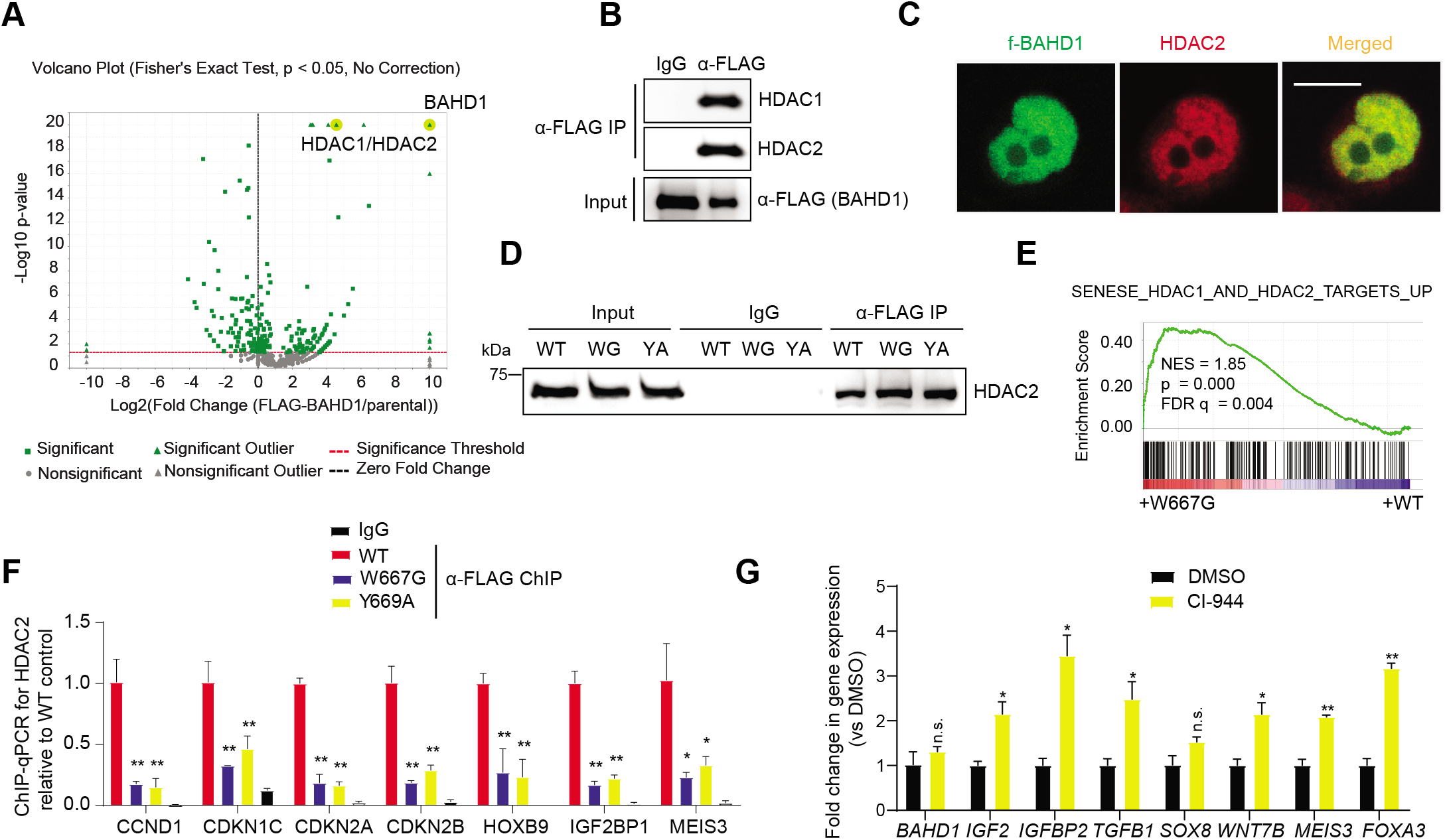
BAHD1 recruits HDACs, mediating BAHD1 target gene repression. **(A)** Volcano plot, based on quantitative mass spectrometry data, demonstrated the enrichment score and statistical significance of the BAHD1 interactors identified from FLAG pulldown of the f-BAHD1 stable expression cells, compared with that of parental HEK293 cells. The x-axis and y axis represent fold-change value (f-BAHD1/parental) and −log_10_ (*P* value) showing statistical significance, respectively. (**B-C**) CoIP (**B**) and IF (**C**) showing both interaction and co-localization of endogenous HDACs with f-BAHD1 in HEK293 cells. (**D**) CoIP for HDAC2 interaction with f-BAHD1, either WT or BAH-mutated, in HEK293 cells. WG and YA indicate W667G and Y669A, respectively. (**E**) GSEA revealing a positive correlation between the W667G mutation of BAHD1 and reactivation of the indicated HDAC target signature. (**F**) ChIP-qPCR for detecting HDAC2 binding to the indicated BAHD1 gene target in HEK293 cells with BAHD1^W667G^ or BAHD1^Y669A^, compared to WT BAHD1. (**G**) RT-qPCR for the indicated BAHD1 target gene in HEK293 cells treated with 5uM of CI-994, relative to DMSO control, for 5 days. Paired t-test was used (n.s., *P* >0.05; *, *P* < 0.05; **, *P* < 0.01).

Next, we aimed to define chromatin changes caused by the disrupted interaction between BAHD1^BAH^ and H3K27me3. Towards this end, ChIP-seq was performed for both gene-activation-related histone acetylation (H3K27ac and H3K9ac) and gene-repression-related histone methylation (H3K27me3 and H3K9me3) in the BAHD1-depleted HEK293 cells that were rescued with exogenous BAHD1, WT or BAHD1^W667G^ (Supplementary Figure 5A). At the BAHD1/H3K27me3 co-targeted genes, overall H3K27ac was found increased in cells expressing BAHD1^W667G^, compared to WT, whereas overall H3K27me3 at these regions was markedly decreased (Figure 6A-B). Meanwhile, while H3K9ac showed a mild increase (Figure 6A-B), H3K9me3 was found to be generally non-existing or very low at these BAHD1-targeted genes, especially TSS-proximal regions (Figure 6A, the bottom panel). Using global ChIP-seq profiles, we also observed a closer correlation between BAHD1 and H3K27me3, in comparison to H3K9me3 (Supplementary Figure 5B). The remodeling of histone modifications (i.e., gain of H3K9ac/K27ac and concurrent reduction of H3K27me3), as well as a lack of H3K9me3, at the BAHD1-bound genomic sites are more evident at a set of BAHD1-repressed genes such as IGF2, TGFB1, SOX8, FOXB1 and CCND1 (Figure 6C and Supplementary Figure 5C). These observations also suggest that H3K27me3, rather than H3K9me3, is more directly involved in targeting BAHD1 to these Polycomb-targeted loci, in contrast to the previous working model (15,18). ChIP-qPCR further verified the increase of histone acetylation at promoters of several BAHD1 target genes (such as IGF2), but not a control house-keeping gene (GAPDH), in cells with BAHD1^W667G^ versus WT (Supplementary Figure 5D). Collectively, we demonstrate that direct binding of H3K27me3 by BAHD1^BAH^ is essential for maintaining a repressive state of histone deacetylation at its Polycomb gene targets.

**Figure 6.**
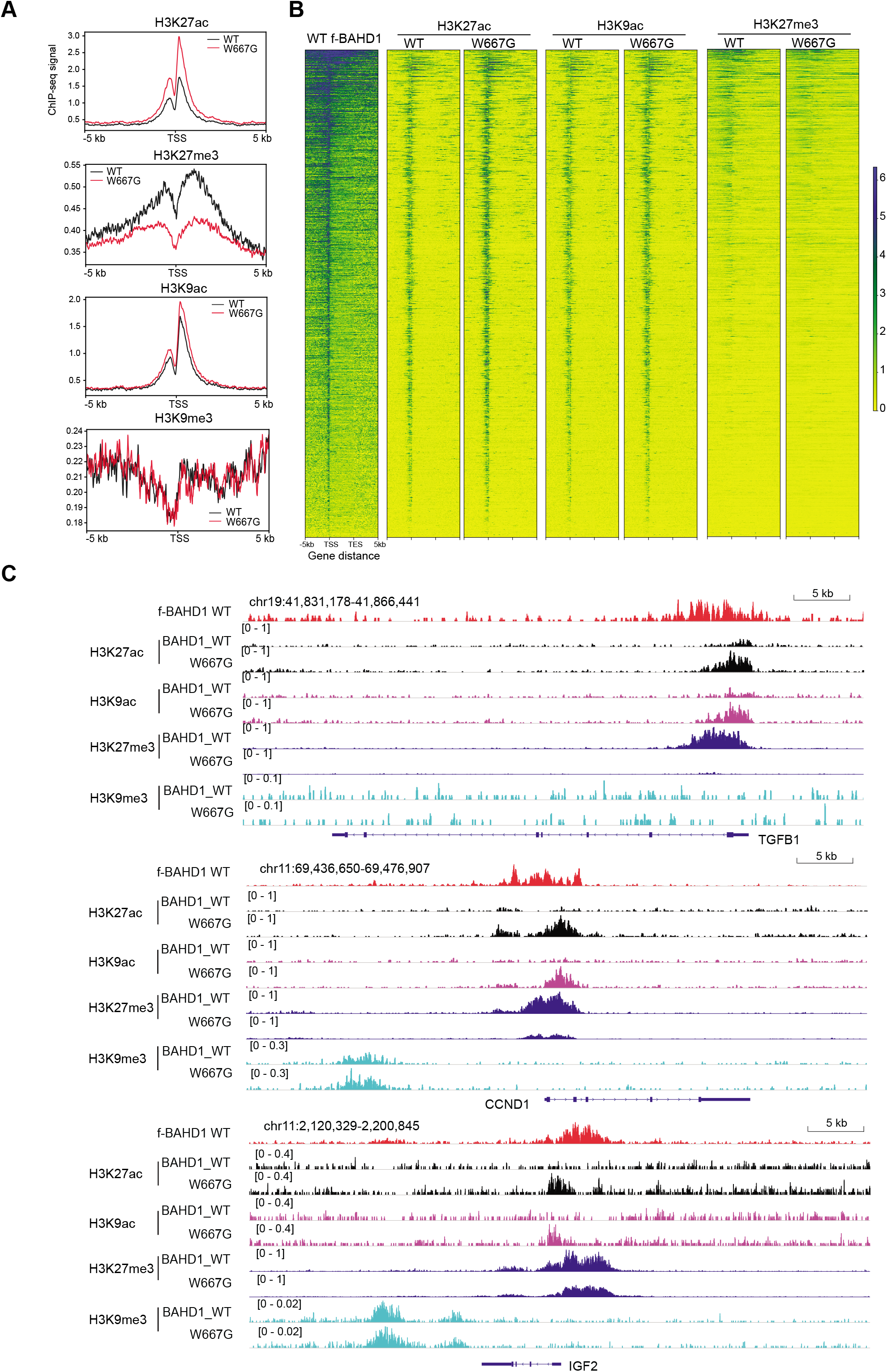
BAHD1 binding to H3K27me3 is required for maintaining histone deacetylation at Polycomb gene targets. **(A-B)** Averaged ChIP-seq signals **(A)** and heatmap **(B)** of H3K27ac, H3K27me3, H3K9ac and H3K9me3 over ±5 kb from TSS to TES of the BAHD1-targeted Polycomb genes in HEK293 cells with expression of either WT or W667G-mutated BAHD1. ChIP-seq reads were normalized to input and sequencing depth. TSS, transcription start site; TES, transcription end site. Color bar, log2(ChIP/Input). **(C)** CUT&RUN signals of WT BAHD1 (FLAG; top) and ChIP-seq profiles of H3K27ac (black), H3K9ac (purple), H3K27me3 (dark blue) and H3K9me3 (light blue) at the indicated Polycomb target in HEK293 cells with expression of either WT or W667G-mutated BAHD1.

### A germline mutation of *Bahd1^W659G^* causes severe neonatal lethality in mice

BAHD1 is essential for animal development. Its global knockout (KO) in mice led to defects in placental and fetal growth (18) whereas the *Bahd1*+/- heterozygous mice showed an anxiety-like behavior (45), indicating a pleiotropic function of BAHD1 among various tissue types and cell lineages. To further assess potential involvement of the BAHD1^BAH^:H3K27me3 interaction during normal development, we generated mice harboring a germline mutation of *Bahd1^W659G^* (Supplementary Figure 6A), equivalent to human *BAHD1^W667G^*. Breeding of the *Bahd1^W659G^* heterozygous mice produced only 2.7% (2 out of a total of 74) of the *Bahd1^W659G^* homozygous pups at birth (Figure 7A), which is significantly lower than the expected frequency among both male and female pups (Figure 7B). Additionally, a few of the surviving *Bahd1^W659G^* homozygous mice exhibited a dwarfism phenotype, compared to the matched WT or heterozygous littermates (Figure 7C-D and Supplementary Figure 6B). Using such a murine model, we further established primary MEF cells that carry either WT *Bahd1* or *Bahd1^W659G^* homozygous alleles. The subsequent RNA-seq profiling revealed the significant enrichment of Polycomb and H3K27me3 target genes among transcripts upregulated in MEFs with Bahd1^W659G^, relative to WT (Figure 7E-F, Supplementary Figure 6C-D, and Supplementary Table 5), which is in agreement with what was observed in the HEK293 cell model. These results thus demonstrate a requirement of the BAHD1^BAH^-mediated H3K27me3 interaction for early embryogenesis and appropriate postnatal development, which was previously unreported.

**Figure 7.**
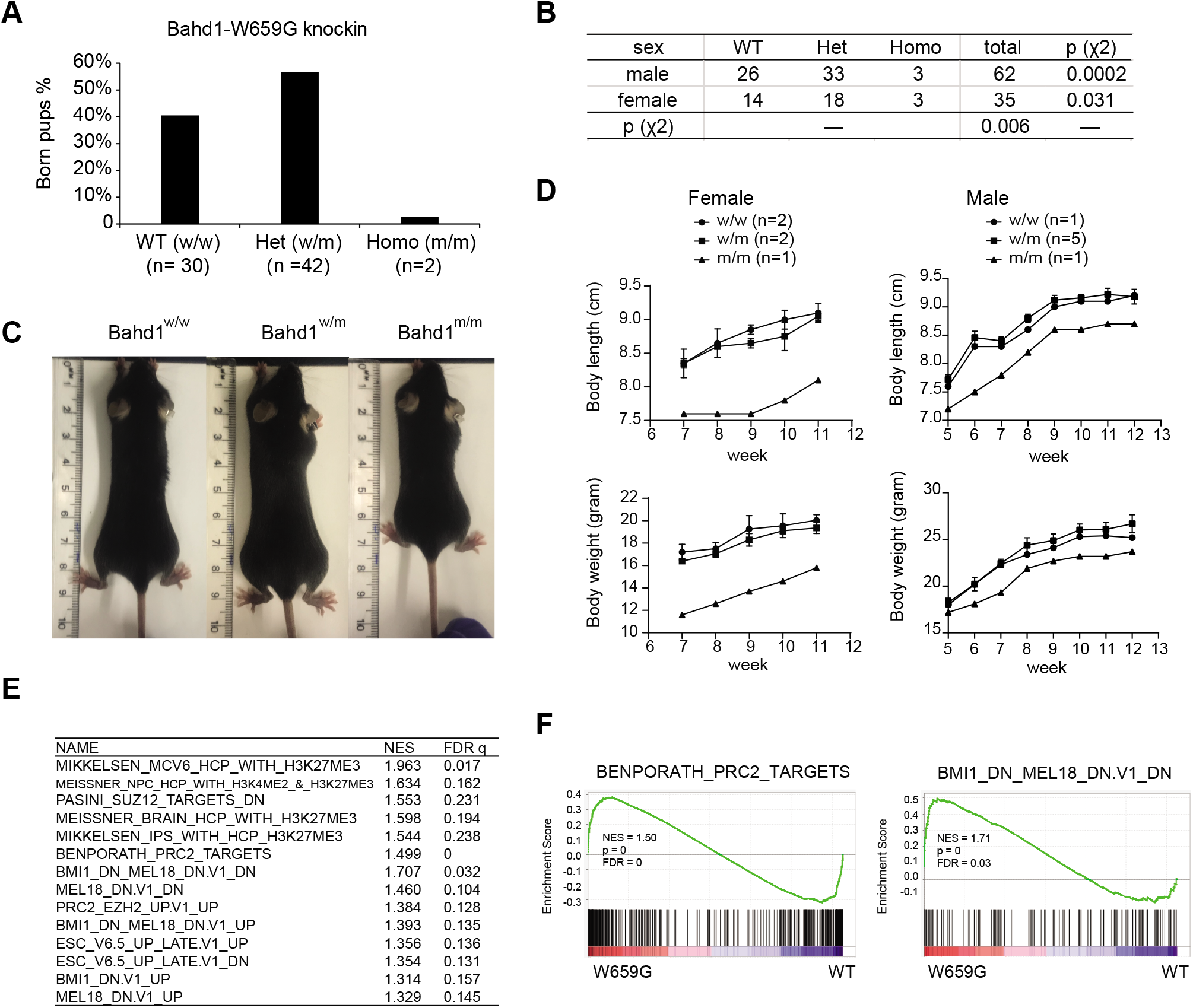
Engagement of H3K27me3 by BAHD1^BAH^ is essential for normal development in mice. **(A**) Bar-chart showing the ratio of newly born pups, which were either *Bahd1* WT (w/w; n= 30) or heterozygous (w/m; n= 42) or homozygous (m/m; n= 2) for *Bahd1^W659G^*, from breeding of heterozygous pairs. **(B**) Chi-square test of the genotypes and sex distribution of the indicated genotype among newly born pups from breeding of the *Bahd1^W659G^* heterozygous pairs. Note that the female pups or the *Bahd1^W659G^* homozygous mutant pups were found to be significantly under-represented. **(C**) Images showing body size of the littermates with the indicated *Bahd1^W659G^* genotype eight weeks post-birth. **(D)** Measurement of body size (top) and body weight (bottom) at the indicated time point post-birth showing that the *Bahd1^W659G^* homozygous mice (female or male; n=1 and 1) were smaller than its heterozygous (n=2 and 3) or WT (n=2 and 5) littermates. (**E-F**) Summary (**E**) and examples (**F**) of GSEA revealed reactivation of the indicated Polycomb gene signature in MEF cells carrying the *Bahd1^W659G^* homozygous mutation, relative to WT.

## DISCUSSION

It was postulated that chromatin modification such as H3K27me3 serves as a context-dependent docking site for recruitment of the so-called reader/effector and associated protein complexes, which impart their respective gene- and genome-regulatory effect (5,46,47). While it is viewed that mammalian cPRC1 binds to H3K27me3 for mediating Polycomb gene silencing, such a pathway is generally absent from the plant and fungus (19,20,22,48). A flurry of recent studies have collectively demonstrated that H3K27me3 is also recognized and directly bound by a set of BAH modules that exist among various organisms covering from Arabidopsis thaliana (19–21,23,24), animal (15–17) and fungus (Neurospora crassa) (22), suggesting a biological significance of the BAH-based functional transduction of H3K27me3. Here, we reported the crucial role for BAHD1^BAH^ in the mammalian Polycomb gene silencing and animal development, which was not explored before. Indeed, although BAHD1^BAH^ was shown to bind H3K27me3 directly (16), it was previously proposed that BAHD1 acts as a multifunctional “scaffold” and its gene-repressive effect involves its associations with H3K9me3 readers and/or other related corepressors (15,18). Through extensive genomic profiling (ChIP-seq, CUT&RUN and RNA-seq) and BAHD1^BAH^ mutagenesis (using 293 cells and a mutation knockin murine model and derived MEF cells), we demonstrate that (i) an intact H3K27me3-engaging module (BAHD1^BAH^) is required for overall BAHD1 binding onto chromatin targets (as shown by chromatin fractionation/association assay and CUT&RUN of BAHD1, WT versus BAH-mutated), (ii) that the direct interaction between BAHD1^BAH^ and H3K27me3 is essential for silencing of a subset of the H3K27me3-demarcated Polycomb targets, which include developmental transcription factors (such as FOXB1 and MEIS3) and imprinted gene (such as IGF2); (iii) that the direct H3K27me3 binding by BAHD1^BAH^ is also required for maintaining histone deacetylation at its Polycomb targets, in agreement with the observed BAHD1:HDAC interaction; and lastly, (iv) that abolishing the BAHD1^BAH^:H3K27me3 engagement causes severe developmental defects in mice such as embryonic lethality and dwarfism, consistent to the known requirement of Polycomb factors for organismal development and cell lineage specification. Notably, the observed chromatin remodeling due to mutation of BAHD1^BAH^ includes increase of H3K27ac/H3K9ac and decrease of H3K27me3. For the latter, it is known that H3K27ac directly antagonizes the PRC2-mediated methylation of the same site and that, following gene reactivation, the productive transcriptional elongation leads to accumulation of the SETD2-mediated H3K36me3, which also suppresses H3K27me3 formation in cis (49).

A previously proposed model emphasizes a multifaceted interaction of BAHD1 with the H3K9me3-specific readers such as HP1 or TRIM28/KAP1 and other repressive partners such as the H3K9 methyltransferase (15,18). Here, we observed that H3K9me3 is generally absent or low at the BAHD1-bound genomic sites, especially TSS-proximal regions (see the bottom panel of Figure 6A and examples in Figure 6C and Supplementary Figure 5C). Based on these results and various phenotypes caused by the H3K27me3-binding-defective mutation of BAHD1^BAH^, we thus favor a view that the direct binding of H3K27me3 by BAHD1^BAH^ provides a primary mechanism for tethering BAHD1 onto its Polycomb gene targets. Besides the H3K27me3-binding BAH motif, BAHD1 can use a N-terminal region as an interface for protein-protein interaction (50) and may form interactions with other gene-repressive pathways of H3K9me3 (15) or DNA methylation (51). Please note that the most frequent and consistent hits of BAHD1 interactors, as identified previously (18) and from this study (Supplementary Table 4), include at least MIER3, MIER1, HDAC1, HDAC2 and TRIM28/KAP1 (in a relatively less frequency), indicating that this set of cofactors form a more stable “core” complex with BAHD1. Please also note that we detected existence of H3K9me3 several to dozens of Kb away from a subset of those BAHD1-bound regions (as exemplified by IGF2, FOXB1 and CCND1 shown in Figure 6C and Supplementary Figure 5C); therefore, putative involvement of TRIM28/KAP1, a BAHD1 partner and H3K9me3-specific reader, is possible at certain BAHD1 targets— for example, it is conceivable that, following initial tethering of BAHD1 to the TSS-proximal regions marked by H3K27me3, a long-range looping interaction between TSS and the distal H3K9me3-demarcated region may form, due to physical association between BAHD1 and the H3K9me3-bound TRIM28/KAP1 (15,18), which possibly mediates further stabilization and/or spreading of the BAHD1 complex on chromatin. While further studies are warranted to delineate the molecular details, the BAHD1-induced heterochromatin formation is likely to be context-dependent (such as chromatin environment and interactors) and involves a multilevel mechanism, of which the direct binding of H3K27me3 by BAHD1^BAH^ represents one of the critical, possibly primary, events.

A significant finding of this report lies in a clear demonstration of BAHD1^BAH^ as a functional H3K27me3 reader in the mammalian cells, which acts to maintain appropriate Polycomb gene-silencing program for sustaining normal development. Considering the existence of cPRC1 and recent identification of BAHCC1^BAH^ as H3K27me3-specific readers (17), the chromodomain (cPRC1)- and BAH-mediated functional readout of H3K27me3 can coexist in mammalian cells, imparting their respective gene-silencing effects through distinctive molecular pathways. Besides a catalytic function for inducing H2AK119ub1, cPRC1 induces chromatin compaction via cofactor-mediated phase separation and condensation formation (12–14). On the other hand, BAHCC1 (17) and BAHD1 both interact with HDACs, thus linking H3K27me3 together with histone deacetylation, an integral step of gene repression. Conceivably, the chromodomain- and BAH-containing effectors of H3K27me3 may potentially cooperate, synergize and/or compete, the detail of which awaits additional analysis. It is also possible that, due to differential expression patterns, their overall contributions to Polycomb gene silencing vary among various cell lineages and tissue types. The mRNA levels of Bahd1 and Bahcc1 are generally lower than those of cPRC1 genes in embryonic stem cells (data not shown) whereas their expression can be more comparable in other cell types such as 293 cells. Overall, the BAH module–based transduction of H3K27me3 is not only functionally important but also evolutionarily ancient, existing in the fungus, animal and plant.

Over dozens of the chromatin-associated proteins from different organisms carry a BAH motif (52). It is noteworthy to mention that these different BAH domains demonstrate functional divergence. In contrast to a set of the H3K27me3-“reading” BAHs mentioned above in fungus, plant and animal, the BAH domain of ORC1 preferentially recognizes histone H4 lysine 20 dimethylation (H4K20me2) in metazoan species (53). In maize, a methyltransferase termed ZMET2 uses its BAH domain to recognize and bind H3K9me2, thereby establishing a crosstalk between the repressive histone modification and DNA methylation (54). In yeast, the BAH motif of Rsc2 was shown to bind recombinant H3 in vitro whereas that of Sir3 contacts with nucleosome to induce heterochromatin formation (55). The molecular basis underlying the observed functional divergence among BAH modules remains to be fully characterized. Importantly, deregulation of the BAH-containing proteins is often associated with pathogenesis. For instance, we reported that germ-line mutation of either Bahcc1^BAH^ (17) or Bahd1^BAH^ in mice caused embryonic lethality and dwarfism. The germ-line mutation of ORC1’s H4K20me2-binding BAH domain is responsible for Meier-Gorlin syndrome, a form of primordial dwarfism (53,56). Somatic mutation at the BAH domains of Polybromo-1 was detected in clear cell renal carcinoma (57). Altogether, BAH evolves as a versatile module family critically involved in various biological processes, which merits further studies.

## DATA AVAILABILITY

All data needed to evaluate the conclusions in the paper are present in the paper and/or the Supplementary Materials. Additional data related to this paper may be requested from the authors. Raw and processed NGS data have been deposited with the National Center for Biotechnology Information (NCBI) Gene Expression Omnibus (GEO) under accession number GSE160354.

## SUPPLEMENTARY DATA STATEMENT

Supplementary Data are available at NAR Online.

## ACKNOWLEDGMENTS

We graciously thank H. Bierne for providing the reagent and the Wang Laboratory members, T. Ptacek, and J. Simon for helpful discussion and technical support. We thank UNC’s facilities, including Imaging Core, High-throughput Sequencing Facility (HTSF), Bioinformatics Core, Animal Models Core Facility and Tissue Culture Facility for their professional assistance of this work. The cores affiliated to UNC Cancer Center are supported in part by the UNC Lineberger Comprehensive Cancer Center Core Support Grant P30-CA016086.

## Author contributions

H.F., Y.G., A.R.K. and L.C., performed experiments. Y.T., W.G., and Y.G., performed genomic data analysis under direction of G.G.W. A.S., R.E., S.B., S.M., and A.T. conducted and analyzed mass spectrometry experiments. H.F., Y.G., H.L., L.C., A.T. and G.G.W. interpreted the data. G.G.W. conceived the project. H.F., Y.G., and G.G.W. prepared the manuscript with inputs of all other authors.

## Supplementary Material

### Supplementary Tables

**Supplementary Table 1.** RNA-seq identified the differentially expressed genes (DEGs) that showed significant up- or down-regulation in HEK293 cells with knockdown (KD) of endogenous BAHD1 and rescue by exogenous BAH-mutated BAHD1 (W667G), in comparison to wildtype (WT) BAHD1.

**Supplementary Table 2.** List of the BAHD1/H3K27me3 co-bound gene targets that exhibited significant upregulation due to the W667G mutation of BAHD1, relative to WT, in HEK293 cells.

**Supplementary Table 3.** RNA-seq identified the DEGs that showed significant up- or down-regulation post-treatment with 5 μM of UNC1999, relative to mock (DMSO), for 3 days in HEK293 cells.

**Supplementary Table 4.** Summary of proteins identified by mass spectrometry following the FLAG pulldown in HEK293 cells, either parental (as a negative control) or with the FLAG-BAHD1 stable expression.

**Supplementary Table 5.** RNA-seq identified the DEGs that showed significant up- or down-regulation in primary mouse embryonic fibroblast (MEFs) cells carrying the W659G homozygous mutation of BAHD1, relative to WT.

**Supplementary Table 6.** Information of the used reagents including primers, antibodies, and kits.

**Supplementary Figure 1.**
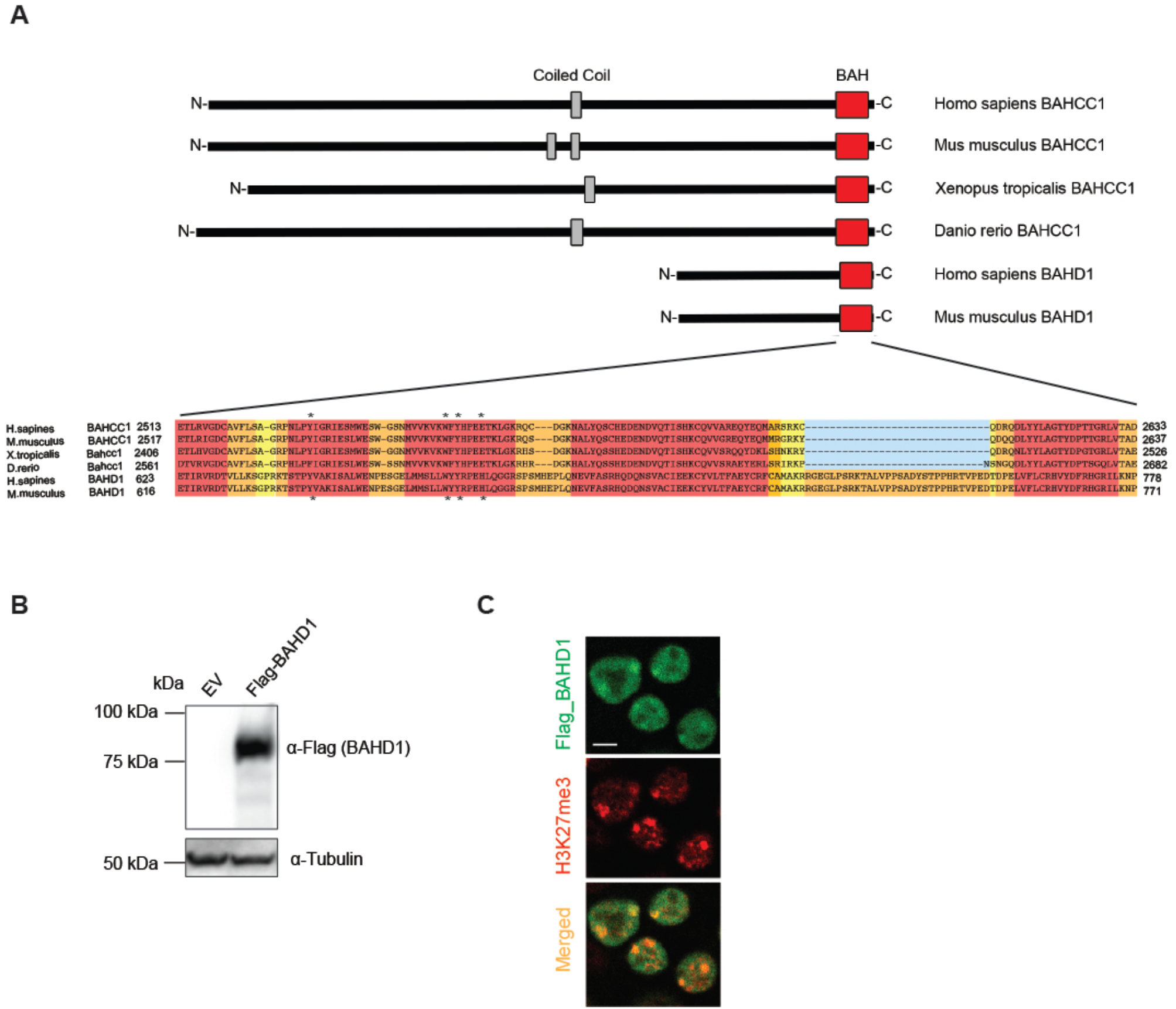
A BAH module of BAHD1 (BAHD1^BAH^) acts as an H3K27me3-specific ‘reader’ in mammalian cells. (**A**) Alignment using the amino acid sequence of the BAH motif embedded in BAHD1 and BAHCC1 of the indicated species. The conserved residues are highlighted in red. The H3K27me3-‘caging’ residues are designated by stars. (**B**) FLAG immunoblotting of the exogenously expressed BAHD1-3xFLAG (detected as a ~85kD protein) post-transfection into HEK293 cells. (**C**) Representative images of confocal immunofluorescence microscopy showing that BAHD1 (green) colocalizes with H3K27me3 (red) in HEK293 cells. Scale bar, 15 μm.

**Supplementary Figure 2.**
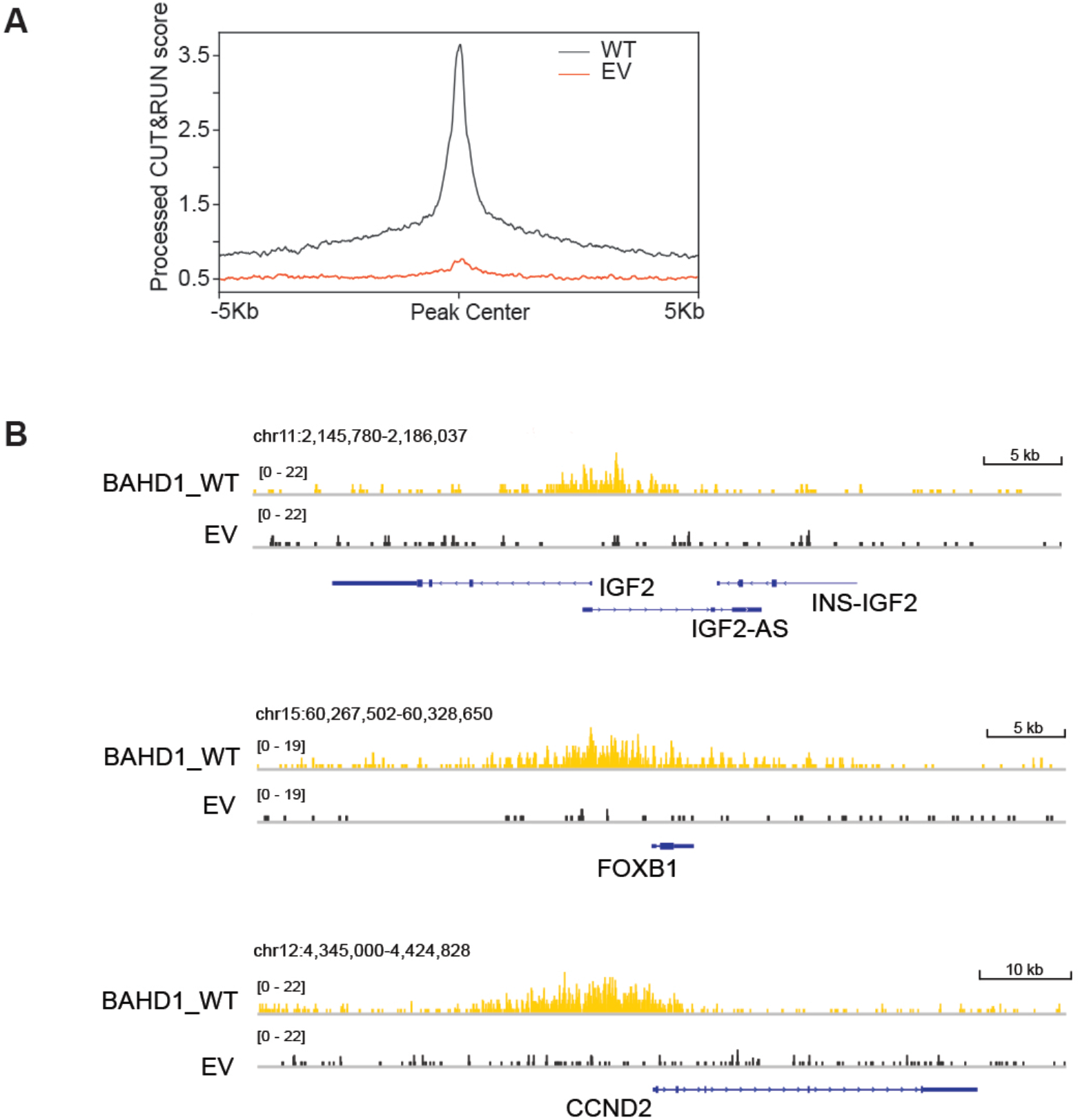
A functional BAHD1^BAH^ motif contributes significantly to BAHD1 binding to the H3K27me3-marked genes in HEK293 cells. **(A)** Averaged FLAG CUT&RUN signals (y-axis) in 293 cells transduced with either empty vector (EV; which serves as a negative control) or WT f-BAHD1. The two samples were processed in parallel and the libraries were subjected to single-end sequencing. The libraries of CUT&RUN samples shown in main Figure 2 were subjected to paired-end sequencing and thus their data analysis were conducted separately. **(B)** IGV views showing the called WT f-BAHD1 peak at the indicated gene in HEK293 cells transduced with EV or WT f-BAHD1.

**Supplementary Figure 3.**
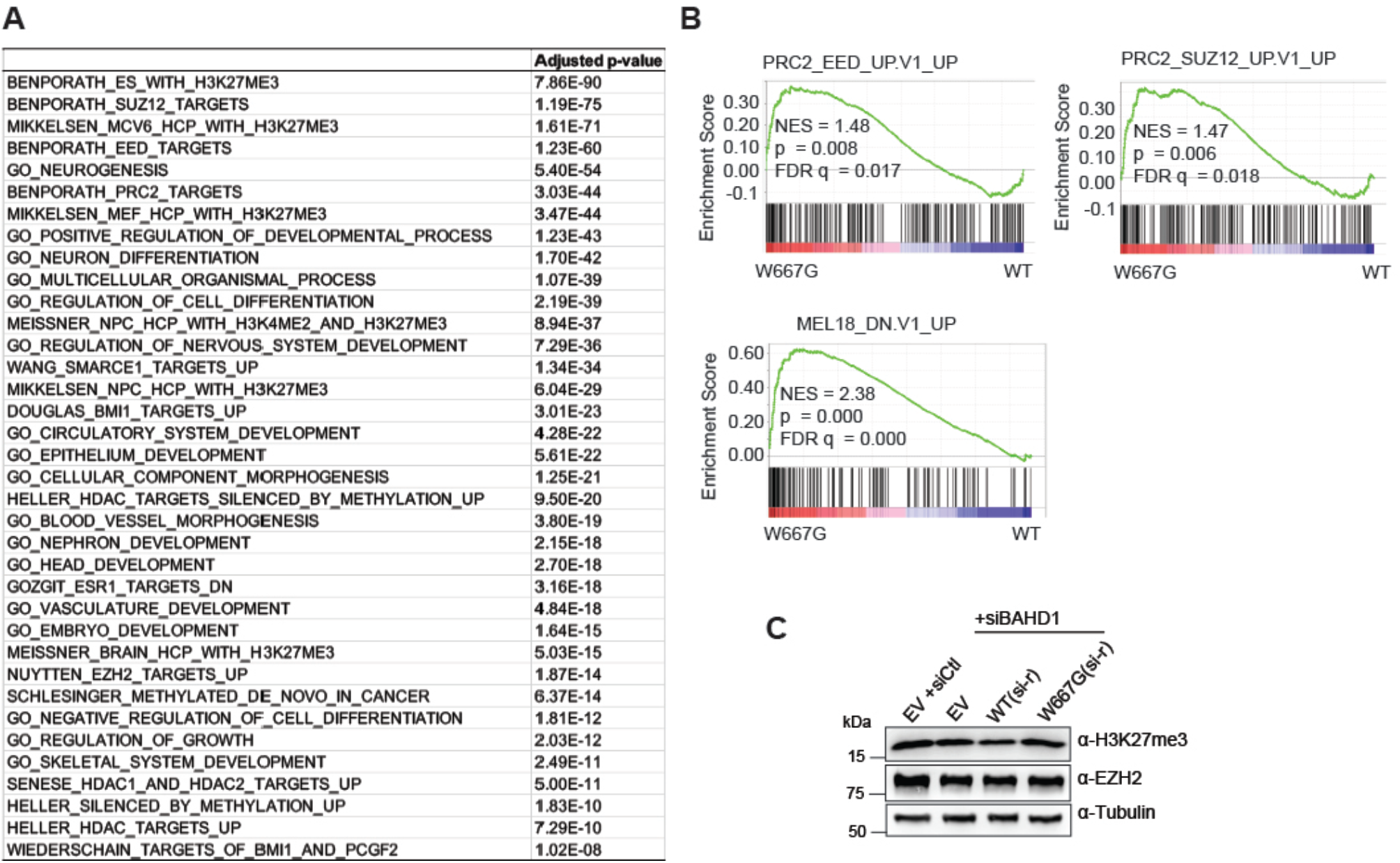
Transcriptomic profiling demonstrates an essential requirement of BAHD1^BAH^ for the BAHD1-mediated Polycomb gene repression in HEK293 cells. **(A)** Platform for Integrative Analysis of Omics data (PIANO) analyses of RNA-seq profiles revealed that, relative to WT, BAHD1^W667G^ is positively associated with derepression of genesets marked by H3K27me3, repressed by PRC2 or PRC1, or related to development. **(B)** GSEA shows that, relative to WT, the H3K27me3-binding-defective mutation of BAHD1^BAH^ is positively correlated to reactivation of gene signatures related to PRC2 or PRC1. **(C)** Immunoblotting for EZH2 and H3K27me3 in the indicated HEK293 cells, which were pre-rescued with either empty vector (EV) or an exogenous siBAHD1-resistant (si-r) BAHD1, WT or BAH-mutated (W667G), followed by transduction of control siRNA (siCtl, lane 1) or siBAHD1 (lanes 2-4) for 48 hours.

**Supplementary Figure 4.**
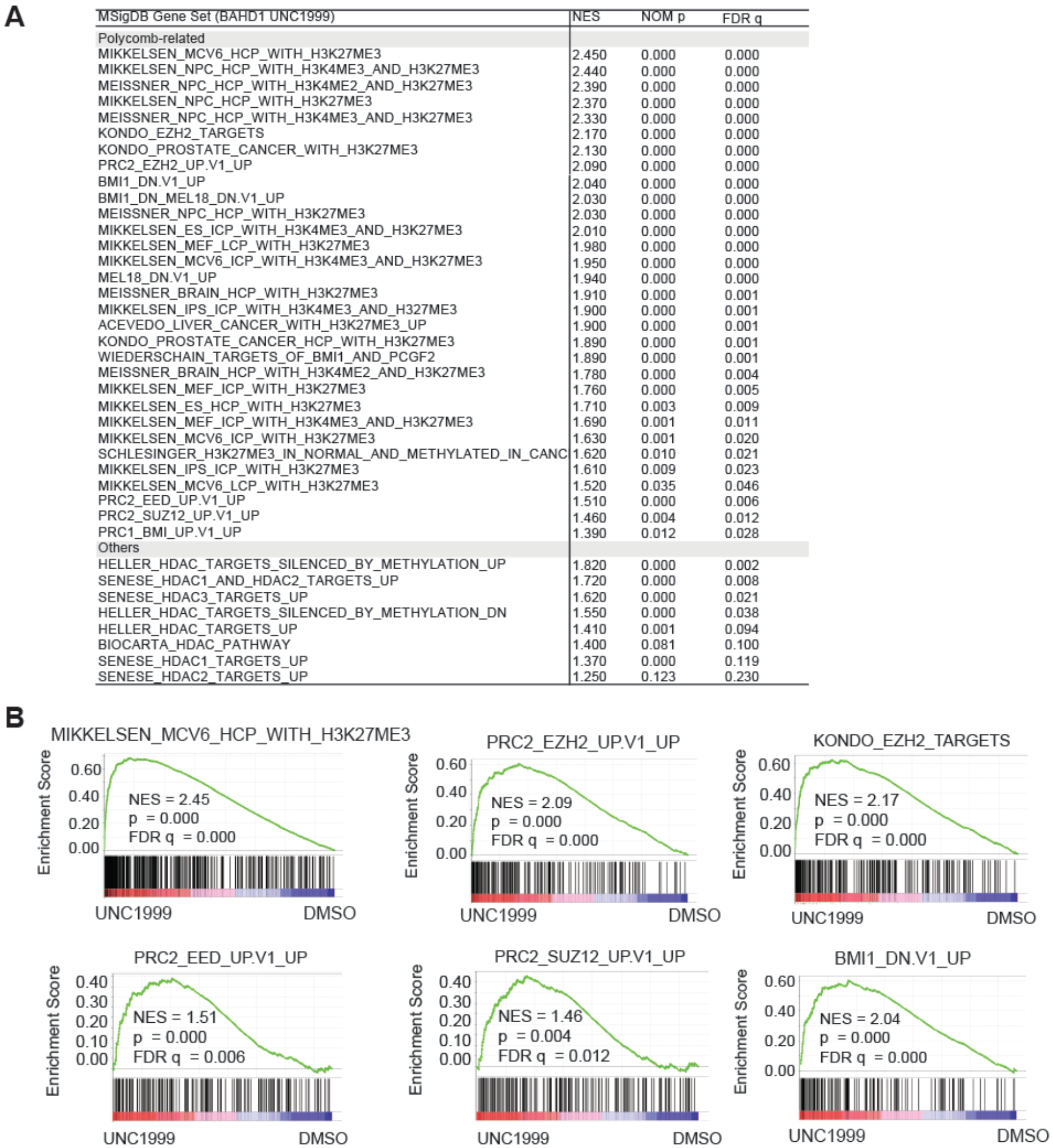
H3K27me3 and BAHD1, which assembles a complex with corepressors such as HDAC1/2, co-repress a set of Polycomb target genes. **(A)** Summary of GSEA using RNA-seq profiles of HEK293 cells post-treatment of UNC1999, relative to DMSO. Representative genesets are categorized into Polycomb (PRC2 and PRC1)-related or others. **(B)** GSEA demonstrates that UNC1999 treatment is correlated with derepression of genesets suppressed by PRC1 or PRC2 or the H3K27me3-bound genes.

**Supplementary Figure 5.**
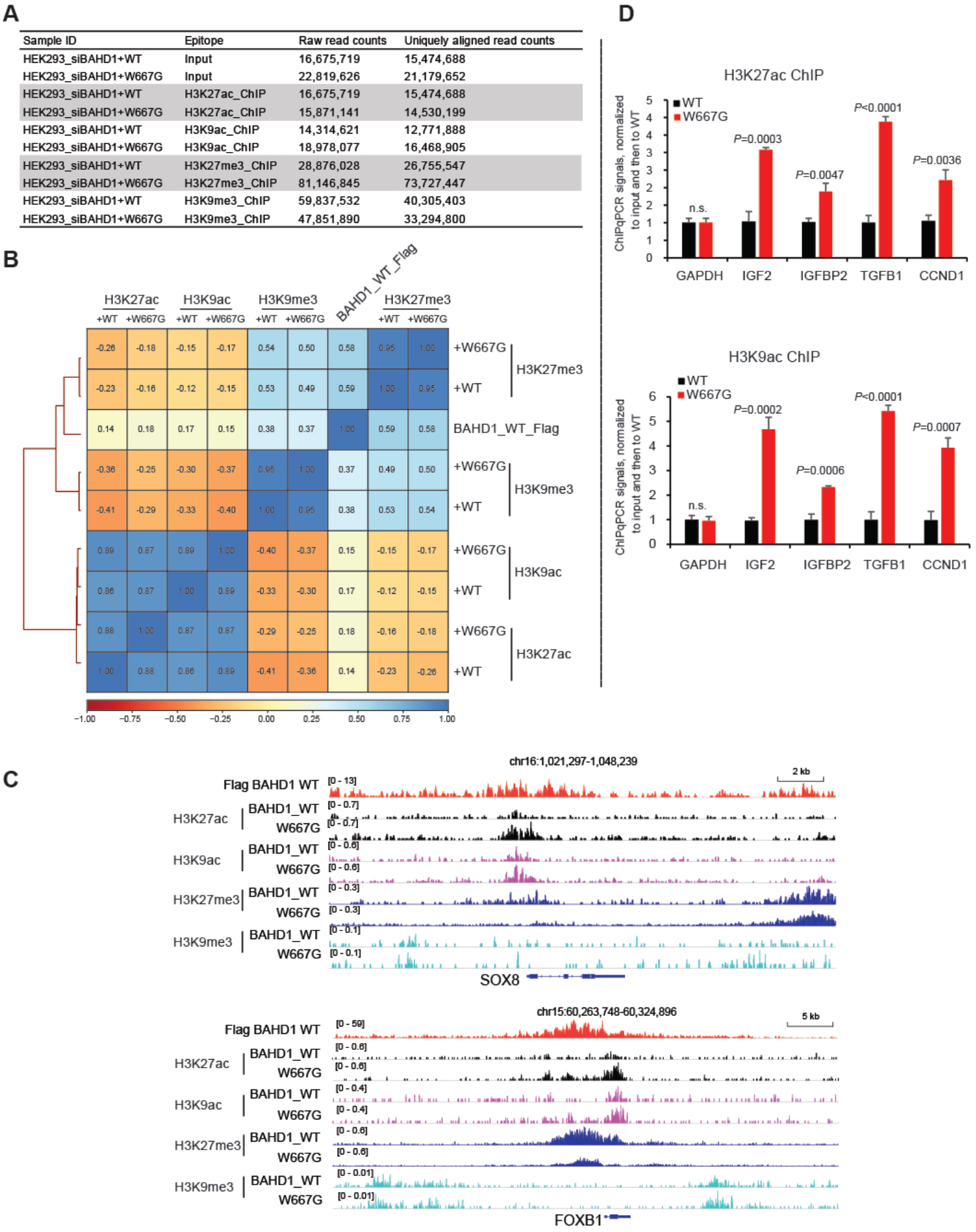
BAHD1 binding to H3K27me3 is required for maintenance of histone deacetylation at Polycomb gene targets. (**A**) Summary of read counts of input, H3K27ac, H3K9ac, H3K27me3 and H3K9me3 ChIP-seq using HEK293 cells (with endogenous BAHD1 depleted) carrying the exogenous WT or W667G-mutated BAHD1. (**B**) Correlation analysis between BAHD1 signals and the indicated histone modification, based on global genomic profiles. (**C**) ChIP-seq profiles of WT BAHD1 (red; FLAG) and H3K27ac (black), H3K9ac (purple), H3K27me3 (dark blue) and H3K9me3 (light blue) at SOX8 and FOXB1 in HEK293 cells (with endogenous BAHD1 depleted) carrying exogenous WT or W667G-mutated BAHD1. (**D**) ChIP-qPCR of H3K27ac and H3K9ac at the indicated gene promoter in HEK293 cells (with endogenous BAHD1 depleted) carrying the exogenous WT or W667G-mutated BAHD1 (n = 3 biologically independent samples). Data are presented as mean ± SD.

**Supplementary Figure 6.**
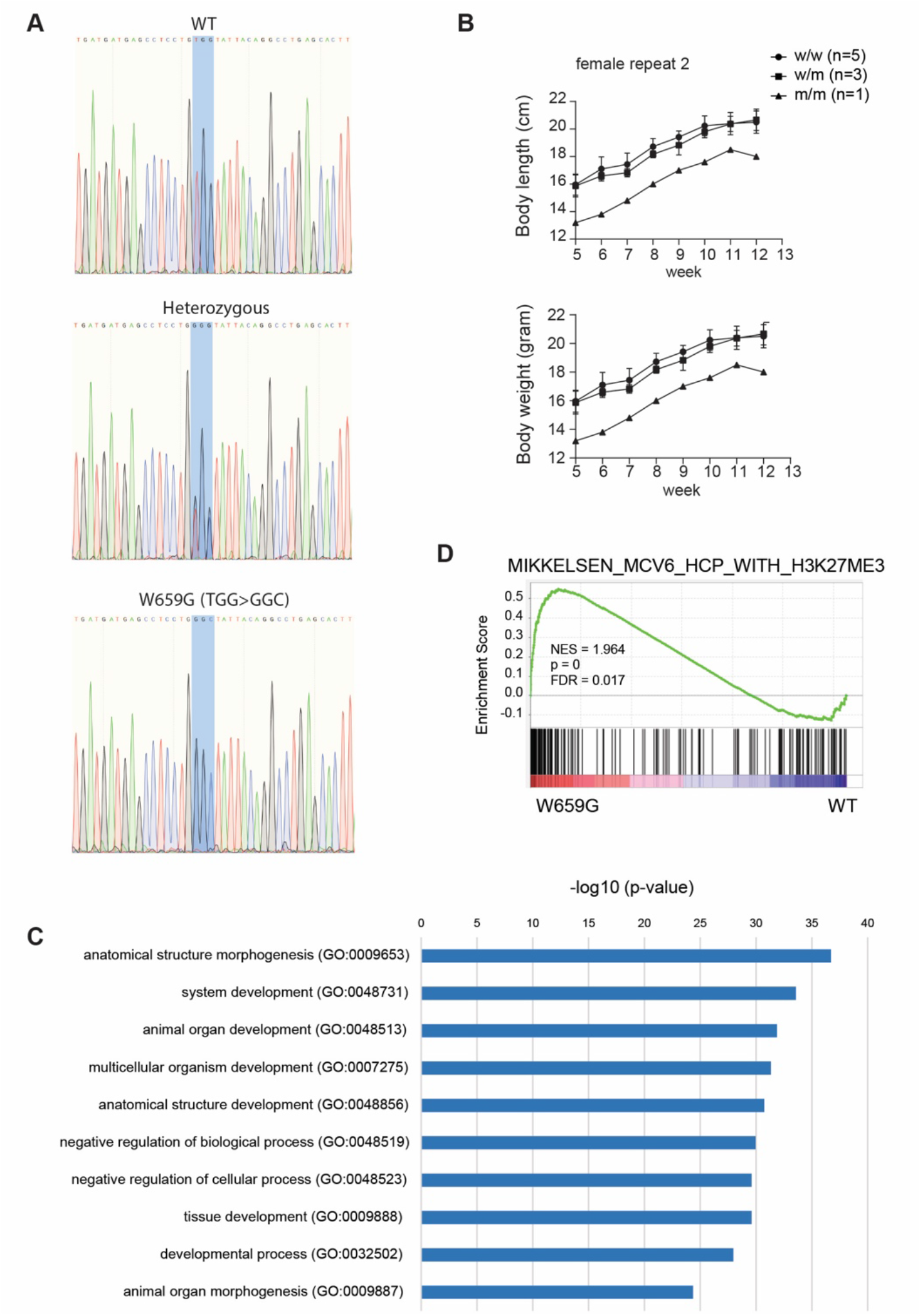
The H3K27me3 engagement by BAHD1^BAH^ is essential for embryonic development in mice. **(A)** Sanger sequencing confirmation of genotype of Bahd1, either WT or with a heterozygous or homozygous W659G (TGG>GGC) mutation, using tissue genomic DNA prepared from mice. **(B)** Measurements of body size and body weight at the indicated time point post-birth showed that a *Bahd1^W659G^* homozygous female mouse (n=1) is smaller relative to its heterozygous (n=5) and WT (n=1) female littermates. (**C**) GO results using the genes upregulated in MEF cells carrying the *Bahd1^W659G^* homozygous mutation, relative to WT. (**D**) GSEA results showed upregulation of the indicated gene in MEF cells carrying the *Bahd1^W659G^* homozygous mutation, relative to WT.

